# Genome-wide reconstruction of rediploidization following autopolyploidization across one hundred million years of salmonid evolution

**DOI:** 10.1101/2021.06.05.447185

**Authors:** Manu Kumar Gundappa, Thu-Hien To, Lars Grønvold, Samuel A.M. Martin, Sigbjørn Lien, Juergen Geist, David Hazlerigg, Simen R. Sandve, Daniel J. Macqueen

## Abstract

The long-term evolutionary impacts of whole genome duplication (WGD) are strongly influenced by the ensuing rediploidization process. Following autopolyploidization, rediploidization involves a transition from tetraploid to diploid meiotic pairing, allowing duplicated genes (ohnologues) to diverge genetically and functionally. Our understanding of autopolyploid rediploidization has been informed by a WGD event ancestral to salmonid fishes, where large genomic regions are characterized by temporally delayed rediploidization, allowing lineage-specific ohnologue sequence divergence in the major salmonid clades. Here, we investigate the long-term outcomes of autopolyploid rediploidization at genome-wide resolution, exploiting a recent ‘explosion’ of salmonid genome assemblies, including a new genome sequence for the huchen (*Hucho hucho*). We developed a genome alignment approach to capture duplicated regions across multiple species, allowing us to create 121,864 phylogenetic trees describing ohnologue divergence across salmonid evolution. Using molecular clock analysis, we show that 61% of the ancestral salmonid genome experienced an initial ‘wave’ of rediploidization in the late Cretaceous (85-106 Mya). This was followed by a period of relative genomic stasis lasting 17-39 My, where much of the genome remained in a tetraploid state. A second rediploidization wave began in the early Eocene and proceeded alongside species diversification, generating predictable patterns of lineage-specific ohnologue divergence, scaling in complexity with the number of speciation events. Finally, using gene set enrichment, gene expression, and codon-based selection analyses, we provide insights into potential functional outcomes of delayed rediploidization. Overall, this study enhances our understanding of delayed autopolyploid rediploidization and has broad implications for future studies of WGD events.

## Introduction

Whole genome duplication (WGD) leading to polyploidy has occurred extensively during eukaryotic evolution (Soltis et al. 2015; Van de Peer et al. 2017). This includes complex WGD histories in plant evolution (Qiao et al. 2019), lineage-defining WGD events ancestral to vertebrates (Simakov et al. 2020) and teleosts (Jaillon et al. 2004), and additional WGDs in several fish families, including salmonids (Lien et al. 2016), cyprinids (Li and Guo 2020) and sturgeons (Du et al. 2020). WGD is widely thought to promote evolutionary diversification through mechanisms that remain incompletely understood (Van de Peer et al. 2017).

Rediploidization follows all WGD events and creates novel genetic diversity (Wolfe 2001). After WGD within the same species (autopolyploidization), rediploidization involves a transition from multivalent (tetraploid inheritance) to bivalent chromosome pairing (diploid inheritance) during meiosis (Furlong and Holland 2002; Lien et al. 2016). Consequently recombination among four alleles ceases, promoting sequence divergence between duplicated genes (ohnologues) residing on distinct chromosomes (Furlong and Holland 2002). This, in turn, creates novel pathways of functional evolution compared to before WGD (Ohno 1970; Conant and Wolfe 2008; Innan and Kondrashov 2010). Rediploidization depends on mutations that promote preferential bivalent pairing during meiosis, such as structural rearrangements (e.g. inversions) and transposable element (TE) insertions (Ohno 1970; Weiss and Maluszynska 2001; Lien et al. 2016). The same rediploidization process will be absent in many allopolyploids (WGD following hybridization of different species) if the consequence is immediate preferential bivalent pairing of the sub-genomes descended from each parent species (Cifuentes et al. 2010; Mason and Wendel 2020), but in theory will occur whenever sequence similarity is sufficient for multivalent pairings to arise, e.g. in segmental allopolyploids (Martin and Holland 2014; Robertson et al. 2017).

A past body of work in salmonid fishes revealed that rediploidization occurred at distinct times in evolution for different genomic regions following an ancestral autopolyploidization (hereafter: ‘Ss4R’) dated at 88-103 Mya (Macqueen and Johnston 2014). The Ss4R is the fourth WGD in salmonid evolutionary history (Berthelot et al. 2014; Lien et al. 2016) following earlier events at the base of vertebrate (Simakov et al. 2020) and teleost evolution (Jaillon et al. 2004). The variable timing of rediploidization for duplicated regions retained from Ss4R is reflected as a ‘snapshot’ within all salmonid genome by distinct levels of sequence divergence among large syntenic blocks of ohnologues (Lien et al. 2016). While rediploidization occurred in the ancestor to all living salmonids in large genomic regions, including several entire chromosomes, speciation occurred before rediploidization was completed in several large genomic segments (Robertson et al. 2017). Consequently, some duplicated chromosome arms share very high sequence similarity (>95%) in all salmonid species (Lien et al. 2016; De-Kayne and Feulner 2018; Campbell et al. 2019; Blumstein et al. 2020) and experienced ohnologue divergence independently in the three salmonid subfamilies^15^, which diverged ~50 Mya (Robertson et al. 2017). This process was previously coined ‘lineage-specific ohnologue resolution’ (‘LORe’) and is thought to be possible whenever the evolutionary transition from multivalent to bivalent pairing (i.e., from tetraploid alleles to ohnologue pairs) occurs after speciation events separating lineages descended from the same WGD event (Martin and Holland 2014; Robertson et al. 2017).

Delayed rediploidization and LORe has implications for phylogenetic inference, as it removes any possibility of 1:1 ortholog relationships among ohnologues retained in affected sister clades, which is the classic expectation following ancestral gene duplication and WGD events (Martin and Holland 2014; Robertson et al. 2017). LORe can readily be mistaken for lineage-specific (e.g., tandem) gene duplication during phylogenetic analyses, if global expectations of WGD (i.e. collinearity among ohnologue pairs on distinct chromosomes) are overlooked (Robertson et al. 2017). When considering the potential evolutionary impacts of WGD events, LORe was hypothesised to promote lineage-specific adaptation (Robertson et al. 2017) and offers a plausible framework to explain frequent observations of time-lags between WGD event and subsequent species or phenotypic diversification regimes (Clark and Donoghue 2017; Carretero‐Paulet and Van de Peer 2020). Following the discovery of delayed rediploidization and LORe in salmonids, many authors have realized the possible importance of these processes for independent WGD events affecting divergent taxa (Macqueen and Johnston 2014; Martin and Holland 2014; Clark and Donoghue 2017; Van de Peer et al. 2017; Rozenfeld et al. 2019; Carretero‐Paulet and Van de Peer 2020). While salmonids represent an outstanding study system, our understanding of rediploidization outcomes in this group of fishes has remained fragmented due to a lack of genome-wide sequence information and/or limited phylogenetic resolution in past reconstructions.

The overarching aim of this study was to reconstruct the post-Ss4R rediploidization process and its outcomes with vastly increased genomic and phylogenetic resolution compared to past work. We sequenced a genome for a species holding a particularly informative phylogenetic position within the salmonid family, and developed a whole genome alignment approach to capture ohnologue regions across genome assemblies generated recently for multiple salmonid species. This unique dataset allowed us to reconstruct genome-wide rediploidization dynamics using phylogenetic methods, capturing two major waves of rediploidization in addition to complex lineage-specific ohnologue divergence histories that scale in complexity with speciation history. Exploiting this new high-resolution ‘map’ of rediploidization, we enhance our understanding of the influence of rediploidization dynamics on a range of gene functional properties. Finally, we discuss the broader implications of our findings for ongoing research into the evolutionary outcomes of WGD events.

## Results

### Reference genome assembly for the huchen (Danube salmon)

To enhance scope to reconstruct rediploidization dynamics across salmonid evolution, we generated a high-quality genome sequence for the huchen (*Hucho hucho*), also known as the Danube salmon (Supplementary Fig 1; Supplementary Tables 1-3; Supplementary Methods 1). This species holds a key phylogenetic position for salmonid comparative genomics. It is part of a species-poor clade within subfamily Salmoninae that is sister to a species-rich clade including Atlantic salmon (*Salmo salar*) and Pacific salmon (*Oncorhynchus*) species (Fig. 1a). The common ancestor of the latter clade is thought to have evolved the capability to migrate into seawater during the life-cycle (termed anadromy), which represents a dominant life-history strategy in extant member species (Alexandrou et al. 2013). In contrast, all species within the huchen’s clade complete the full life-cycle in freshwater, which represents the inferred ancestral state for salmonids (as observed in all grayling species; subfamily Thymallinae) (Alexandrou et al. 2013). Past work has shown that ~25% of the Salmoninae genome experienced rediploidization after the split from Thymallinae (Robertson et al. 2017). Adding the huchen to this study captured the most basal speciation event in Salmoninae, allowing us distinguish regions in the genome that underwent rediploidization in the common Salmoninae ancestor, from regions that experienced lineage-specific rediploidization after the split of huchen from the ancestrally anadromous Salmoninae lineage (Fig. 1a). As the huchen is an endangered species, a reference genome can also be used to support genetic research aimed at wild stock conservation and restoration (Geist et al. 2009; Kucinski et al. 2015).

**Fig. 1.**
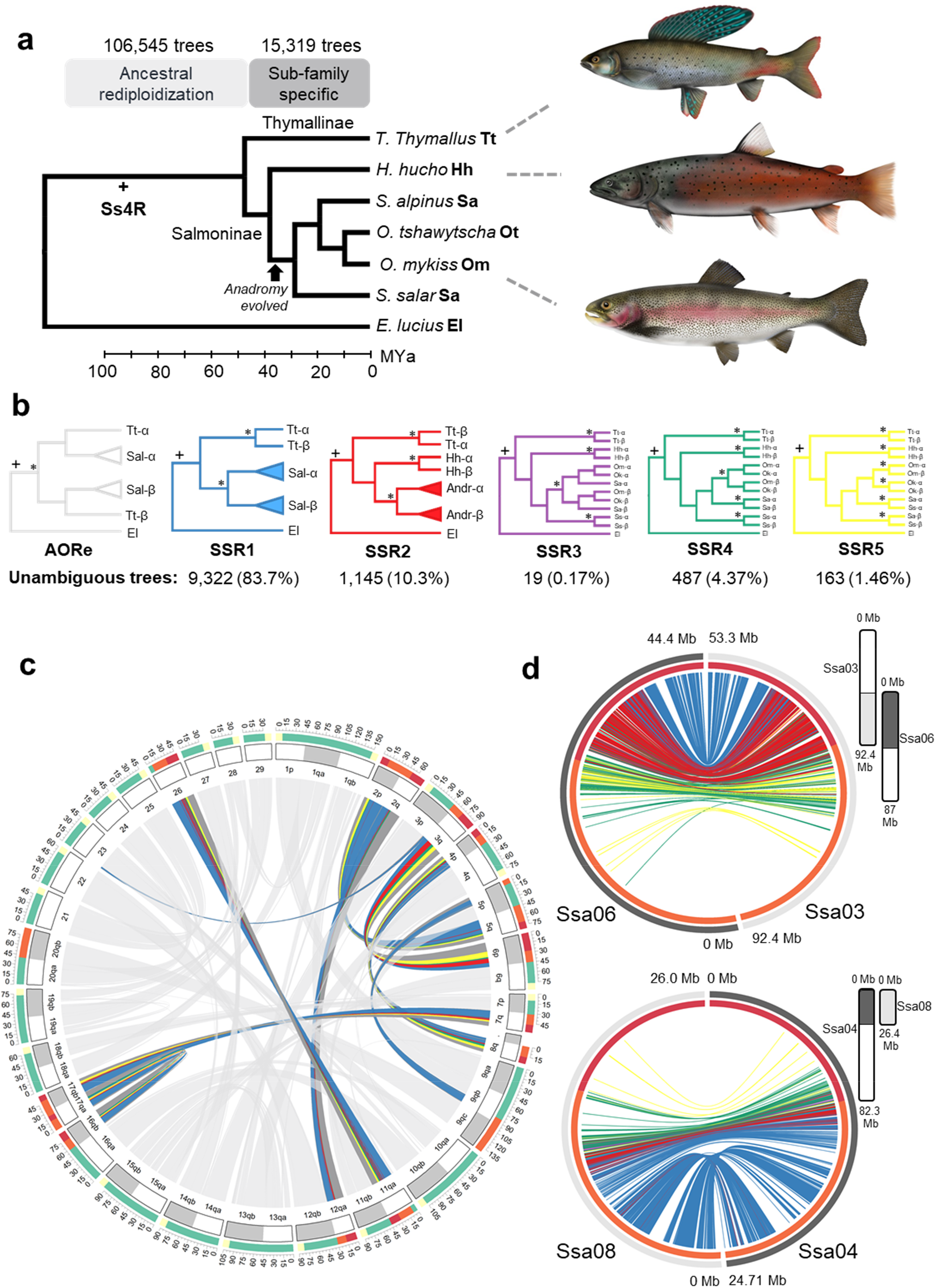
Genome-wide phylogenetic reconstruction of rediploidization history following the Ss4R autopolyploidization. **a**. Phylogeny and divergence times for species used in genome-wide alignment. Also highlighted are the number of captured ohnologue trees in ancestral rediploidization (AORe) and subfamily specific rediploidization (SSR) regions of the genome and the timing of the Ss4R event (Macqueen and Johnston 2014). **b**. The number of phylogenetic trees matching predicted lineage-specific rediploidization scenarios in SSR regions. **c**. Circos plot mapping inferred rediploidization histories (i.e. SSR categories) from ohnologue trees along the Atlantic salmon genome; colours match to the SSR topologies shown in part b. Chromosome arm names follow established nomenclature for Atlantic salmon (Lien et al. 2016). **d**. Circos plots mapping SSR topologies for two example chromosome arms. Additional data provided in Supplementary Figs. 2-10.

Our huchen assembly was generated using Illumina technology from a haploid individual (Supplementary Fig. 1) and had 2.49 Gb total sequence length, contig/scaffold N50 of 37.6/287.3 Kb and 90.2% BUSCO (Simão et al. 2015) completeness (Supplementary Methods 1). An annotated version with 50,114 coding gene models is available on the Ensembl genome browser (https://www.ensembl.org/Hucho_hucho).

### Multispecies genome alignment including ohnologues

With the goal to reconstruct genome-wide rediploidization dynamics in salmonids, we developed a genome alignment approach to capture Ss4R ohnologue regions across multiple species (see Methods). These alignments included species from Salmoninae and Thymallinae as well as northern pike *Esox lucius*, selected as a representative of Esociformes, a sister lineage that diverged before Ss4R (Macqueen and Johnston 2014; Lien et al. 2016; Robertson et al. 2017) (Fig. 1a). We generated multispecies alignments for *a priori* defined syntenic ohnologue blocks retained from Ss4R (Lien et al. 2016) in two genome portions where rediploidization was either ancestral to all salmonids (ancestral ohnologue resolution ‘AORe’ regions; Robertson et al. 2017), or occurred after the split of Salmoninae and Thymallinae (subfamily-specific rediploidization ‘SSR’ regions) (Supplementary Data 1 and 2).

Using this approach, 3,709,704 and 511,436 raw alignments were generated across the ohnologue blocks in AORe and SSR regions, respectively (Supplementary Dataset 3, 4). We next applied a step to filter the alignments according to the number of sequences represented to ensure we retained informative multispecies representation of ohnologue regions (see Methods), and further applied Gblocks (Castresana 2000) to remove low confidence positions in each alignment. This led to 106,545 (sum length: 92.2 Mbp) and 15,319 (sum length: 32.3 Mbp) high-quality alignments spanning the AORe and SSR regions, respectively (Supplementary Data 5, 6), which were used to generate the same number of maximum likelihood phylogenetic trees (Fig. 1a). The alignments and trees used in subsequent analyses are provided and described in Supplementary Data 7 and 8, respectively.

### High-resolution reconstruction of lineage-specific rediploidization histories

Using our genome-wide dataset of phylogenetic trees, we classified rediploidization histories based on the onset of ohnologue divergence (Robertson et al. 2017) in the SSR regions of the genome. Five distinct tree topologies (hereafter: SSR1, 2, 3, 4 and 5) capture the spectrum of predicted rediploidization histories according to the species tree (Fig. 1b). For instance, SSR1 indicates independent rediploidization (i.e., ohnologue divergence) in Thymallinae and the common ancestor of Salmoninae members. At the other end of the spectrum, SSR5 indicates independent rediploidization in every species (Fig. 1c).

11,139 (72.7%) of the available 15,319 trees unambiguously fitted to one of the predicted SSR histories (Supplementary Data 9). Among these, most trees (83.7% of 11,139) supported rediploidization in the Salmoninae ancestor (SSR1 topology; Fig. 1b). 10.3% of the trees matched to expectations of two independent rediploidization histories in Salmoninae, once during *Hucho* evolution and again in the ancestor to *Salmo, Salvelinus* and *Oncorhynchus* (SSR2 topology; Fig. 1b). A smaller number of trees matched to predictions of additional rediploidization events nested within Salmoninae (SSR3-5 topologies; Fig. 1b).

By positionally mapping the recovered phylogenetic topologies along the Atlantic salmon genome (Lien et al. 2016), it was evident that SSR classifications are not randomly distributed, with large genomic regions dominated by common phylogenetic signals (Fig. 1c, d; Supplementary Figs 2-10). Different ohnologue blocks (chromosome arm nomenclature used standard for *S. salar*; e.g. Lien et al. 2016) have distinct rediploidization histories, with most dominated by the SSR1 category. Some regions, including Ssa03-06 and Ssa04-08 (Fig. 1d; Supplementary Fig. 2, 3) in addition to Ssa02-12 (Supplementary Fig. 4) harbour large genomic regions dominated by the SSR2 category. The mapping of SSR topologies was closely associated with the level of sequence divergence between ohnologue regions (Fig. 1c, d). Duplicated regions sharing >97% identity either represent SSR4/SSR5 topologies or more commonly, missing data (Fig. 1d); these regions often harboured insufficient data to pass our alignment filtering criteria (see Methods). In particular, these alignments often contained a single sequence in multiple species, suggesting collapse during genome assembly due to the high similarity of ohnologue sequences (e.g. Varadharajan et al. 2018).

While the genomic location of different SSR tree topologies was strongly clustered, some regions contained a mixture of different, closely related phylogenetic topologies; a pattern prevalent in regions dominated by SSR2-4 trees (e.g. Fig. 1c, d). While this could reflect weak phylogenetic signal, the average filtered alignment length was >2kb, and our quality control efforts removed ambiguous and weakly-supported topologies (see Methods). We initially hypothesised that this observation reflected errors in the reference Atlantic salmon genome, which has lowest accuracy in late rediploidization regions showing high ohnologue similarity (Bertolotti et al. 2020). To test this, we positioned all trees mapping to Ssa03-06 (1,898 trees) along the homologous regions of a more recent long-read based genome for brown trout *S*. *trutta*, with markedly higher contiguity. An identical pattern was observed (Supplementary Fig. 11-12), suggesting assembly error was not an important factor, or that both assemblies suffered the same issue. Another possibility considered was that the mixing of phylogenetic signals resulted from using genome alignment data, rather than gene trees. To test this, we mapped 236 high-quality gene trees including Ss4R ohnologues across the same salmonid species (generated following Bertolotti et al. 2020) to Ssa03-06, and observed the same pattern (Supplementary Fig. 13). Our current interpretation is that the mixing of related SSR topologies is explained by small scale intrachromosomal rearrangements that re-ordered the position of genomic regions sharing common rediploidization histories.

To summarize, these findings reveal complex histories of lineage-specific ohnologue divergence resulting from delayed rediploidization, which scale in number and complexity with the number and timing of speciation events during salmonid evolution.

### Spectrum of rediploidization ages across the genome

The complex lineage-specific rediploidization histories inferred in SSR regions led us to ask if rediploidization timing varied across AORe regions. The challenge to answering this question is that all trees in AORe regions are characterized by the same topology, representing a single onset of ohnologue divergence in the salmonid common ancestor (Macqueen and Johnston 2014; Robertson et al. 2017) (Fig. 1b). In other words, unlike SSR regions, tree topologies provide no information on rediploidization age.

As an alternative approach to empirically estimate rediploidization age, we applied the Bayesian relaxed clock approach MCMCtree (Yang 2007) using the genome-wide alignment data and published temporal constraints on species divergence (Campbell et al. 2013; Macqueen and Johnston 2014; Lecaudey et al. 2018) 04/06/2021 07:46:00to estimate the onset of ohnologue divergence (after Macqueen and Johnston 2014). For AORe regions, we concatenated sequence alignments across 23 defined ohnologue regions in the Atlantic salmon genome (Lien et al. 2016), assuming each represented a single shared rediploidization history (Supplementary Data 10). The estimated rediploidization ages (Bayesian mean) ranged from 68 to 106 Mya (Fig. 2a; Supplementary Data 11). While representing a spectrum of rediploidization ages, the 95% credibility intervals overlapped for 21 out of the 23 regions (Fig. 2a). This makes it is impossible, even with the maximal available sequence data, to distinguish scenarios where rediploidization occurred concomitantly from scenarios where rediploidization was staggered across tens of millions of years. Nonetheless, the duplicated region Ssa09-20 clearly underwent rediploidization later in time, as its upper 95% credibility interval does not overlap with the lower 95% credibility interval for all but one of the remaining AORe regions (Fig. 2a). This analysis suggests that AORe regions representing 61.4% (1.37 out of 2.24 Gb) of the chromosome-anchored Atlantic salmon genome underwent rediploidization no later than ~80 Mya according to the lower 95% credibility intervals (Fig. 2a; Supplementary Data 11). Finally, the most ancient inferred rediploidization ages indicated an older absolute date for the timing of Ss4R than current estimates (Berthelot et al. 2014; Macqueen and Johnston, 2014; Lien et al. 2016), with Ssa07-18 showing a mean rediploidization age of 106 Mya (Fig. 2a; Supplementary Data 11).

**Fig. 2.**
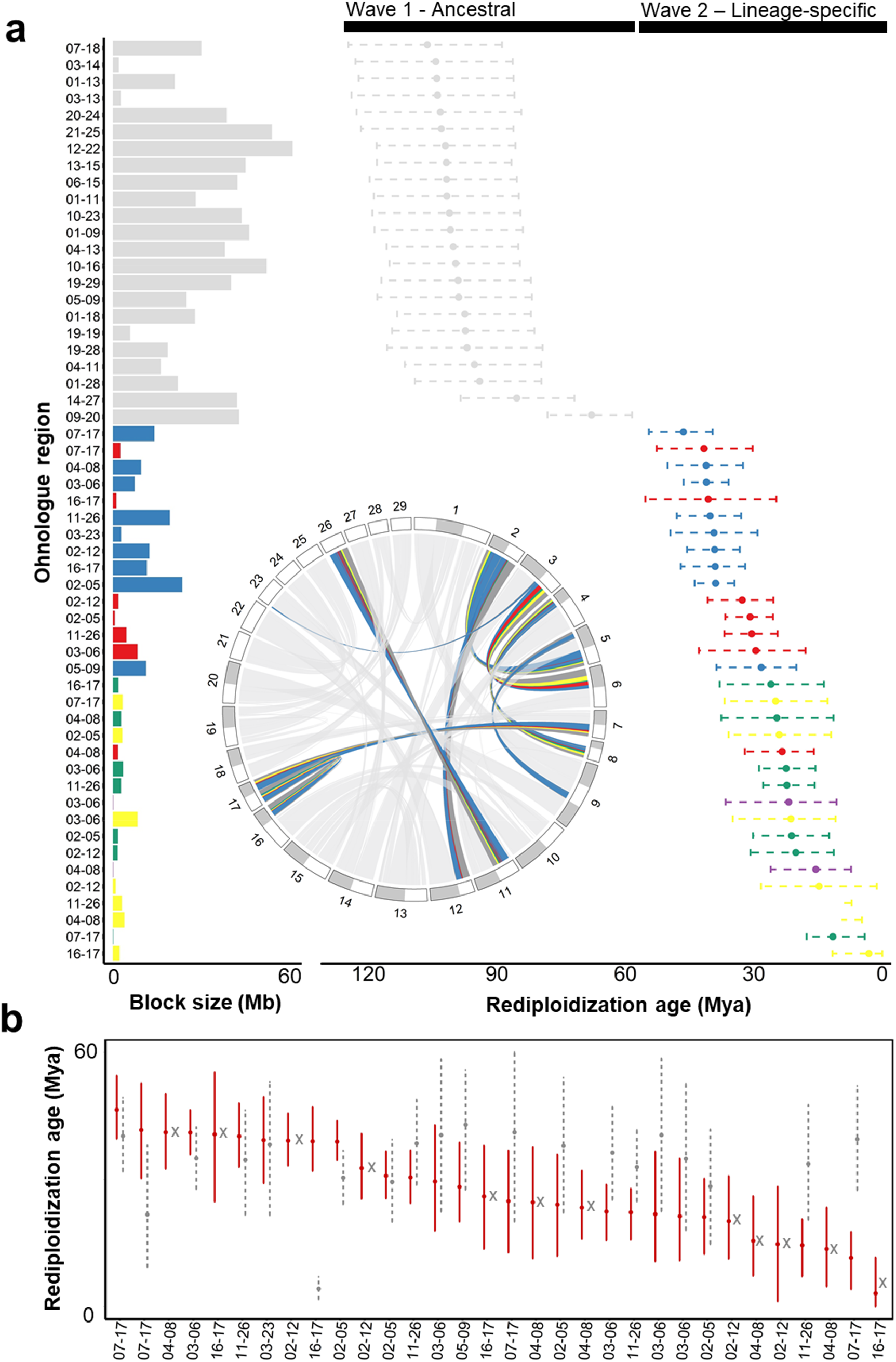
Absolute rediploidization age estimation for ohnologue blocks retained from the Ss4R autopolyploidization. **a**. Onset of ohnologue divergence estimated by MCMCtree using concatenated genome alignments (Supplementary 10). For each ohnologue block, the plotted circle is the posterior mean, and the dotted line is the 95% credibility interval. **b**. Comparison of estimated rediploidization age in SSR regions for Atlantic salmon (red lines) and European grayling (grey lines). Grey crosses indicate regions of presumed assembly collapse in the European grayling genome due to highly delayed rediploidization or maintenance of tetraploidy.

The same approach allowed us to estimate rediploidization ages for SSR regions, where it is possible to infer the timing of lineage-specific ohnologue divergence in different species. This was done for Atlantic salmon and European grayling, which split ~50-60 Mya (Campbell et al. 2013; Macqueen and Johnston 2014; Lecaudey et al. 2018), by concatenating alignments for genomic regions showing strong support for different SSR topologies (Supplementary Data 10; visualized in Fig. 1c). We observed cases where ohnologue regions have similar rediploidization age estimates in both lineages, despite their independent histories of divergence, with overlapping 95% credibility intervals, as well as regions with very different rediploidization ages. For example, a large SSR1 region within Ssa05-09 has the oldest rediploidization age estimate (Bayesian mean: ~43 Mya) in European grayling; ~15 Mya older than in Salmoninae (Bayesian mean: ~28 Mya) (Fig. 2b). Two chromosome arms, Ssa04-08, Ssa02-12, containing regions (spanning SSR1-5 in Salmoninae) with estimated rediploidization ages from 14-42 and 15-39 Mya, respectively, are represented by a single sequence in the European grayling genome, indicative of assembly collapse and potentially maintenance of tetraploidy (Varadharajan et al. 2018). Ssa16-17 had estimated rediploidization ages of 3 to 41 Mya in Salmoninae (spanning SSR1-5), but again showed assembly collapse in the European grayling genome (Fig. 2b).

To summarize, this analysis reveals distinct waves of ancestral and lineage-specific rediploidization following Ss4R, separated by a period of comparative stasis (where just one genomic region underwent rediploidization) spanning 17-39 million years, in addition to the existence of several homologous genomic regions in different salmonid subfamilies with markedly different rediploidization ages.

### Does rediploidization age influence the retention of gene functions?

We next asked if functions encoded by genes across the genome are influenced by rediploidization history using gene set enrichment analysis. The rationale for this question is that different periods in salmonid evolution were coupled to unique genetic and ecological pressures, which may have shaped selection on the retention of gene functions in genomic regions with unique rediploidization histories. For instance, during the period immediately post-Ss4R, rediploidization and ohnologue divergence occurred at a time when molecular adaptations were likely required to recover fundamental cellular and housekeeping functions disrupted by autotetraploidization (Storchová et al. 2006; Hollister 2015; Gillard et al. 2021). Conversely, the more recent period of salmonid evolution, characterized by LORe in large genomic regions, is coupled to species diversification and lineage-specific selective pressures.

Our past work in Atlantic salmon (Robertson et al. 2017), followed by a study in rainbow trout (Campbell et al. 2019) identified differences in gene functional enrichment among sets of Ss4R ohnologues sampled from genomic regions that experienced ancestral (i.e. AORe regions) or delayed rediploidization (i.e. SSR regions). We advanced these efforts by i) taking advantage of our new high definition map of rediploidization history (Fig. 1c), and ii) using an expanded set of Ss4R ohnologue pairs and singleton genes (where one ohnologue in a pair was lost during evolution) defined by Bertolotti et al. 2020. Adding singletons to the analysis allowed us to capture biases in functional enrichment compared to ohnologues, as reported after many WGD events (e.g. Blomme et al. 2006; Smet et al. 2013; Inoue et al. 2015; Han et al. 2016; Parey et al. 2020).

We extracted eight non-overlapping gene sets from the Atlantic salmon genome, representing regions with four distinct rediploidization histories (Fig. 1c): AORe (14,325 ohnologues; 5,887 singletons), SSR1 (3,140 ohnologues; 539 singletons), SSR2 (650 ohnologues; 78 singletons) and SSR3, 4 and 5 combined (hereafter: ‘SSR345’) (426 ohnologues; 162 singletons) (data in Supplementary Data 12 and 13). We tested for enrichment (*P*<0.001) in each gene set using GOslim terms versus the background of all Atlantic salmon genes (Fig. 3; Supplementary Data 12). For standard GO terms, many functions are represented by a small number of genes, which led to a concern that comparing genes extracted from non-overlapping genomic regions with different sizes could cause to substantial biases. GOslim provides a course description of gene functions, typically inclusive of hundreds to thousands of genes per term, which we rationalized would largely circumvent this issue, given that the contributing genes would be represented broadly within the non-overlapping gene sets compared.

**Fig. 3.**
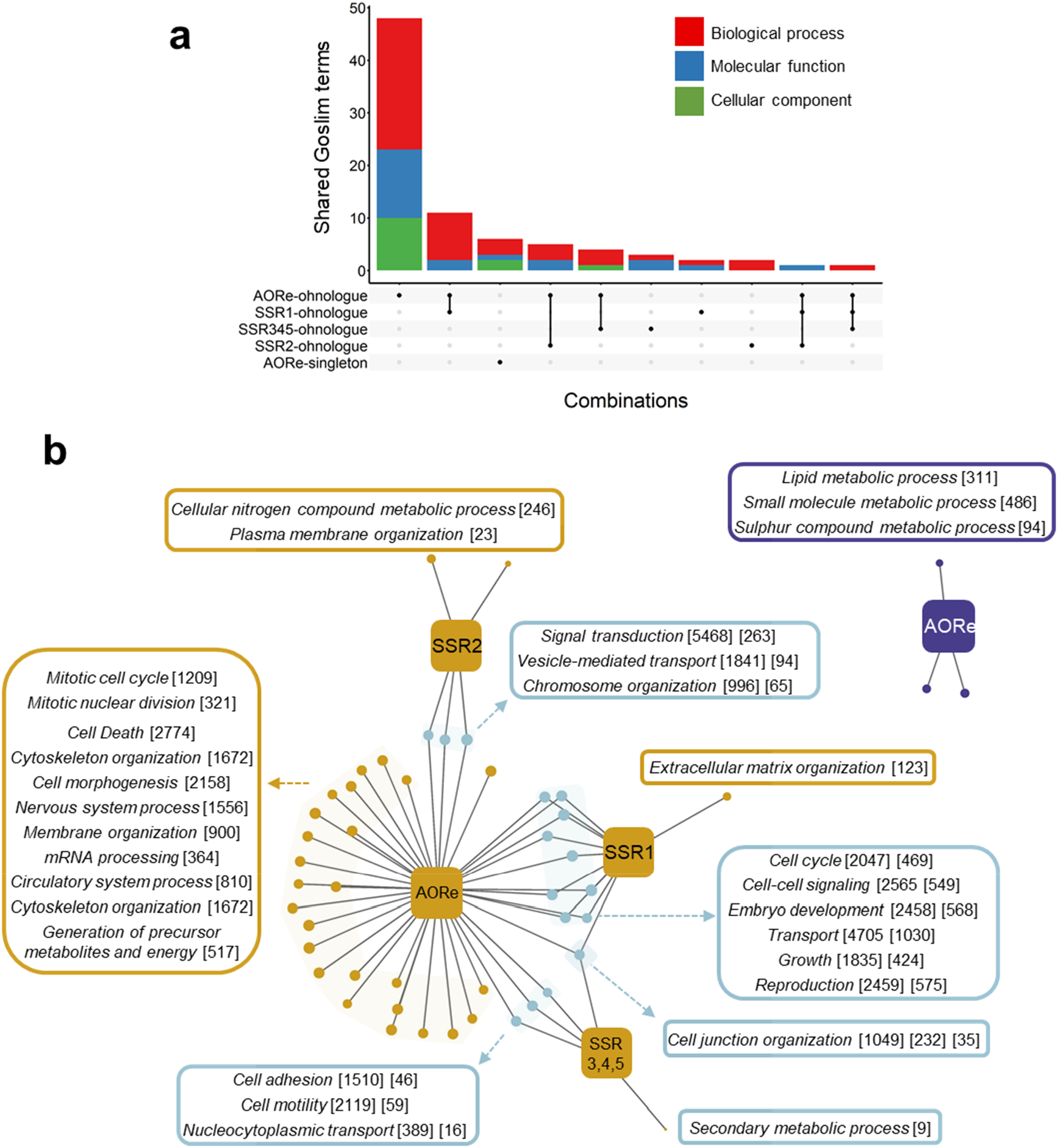
Gene set enrichment analyses contrasting regions with distinct rediploidization ages in the Atlantic salmon genome. **a**. Upset plot highlighting shared enriched GOslim terms among Ss4R ohnologues and singleton genes extracted from AORe and SSR regions. **b**. Network visualizing shared enriched GOslim biological processes among the same categories. Each node represents a unique category (yellow nodes: Ss4R ohnologues; purple nodes: Ss4R singleton genes) in the genome and the lines extending from nodes connect to either category-specific (ending in yellow/purple circles for ohnologues/singletons) or category-shared Goslim terms (ending in turquoise circles). Examples category-specific and category-shared Goslim terms are provided. Full data in Supplementary Data 12.

The AORe ohnologue set, which contained the largest number of genes, showed the largest number of overrepresented GOslim terms, most of which were not shared with other gene sets, including ohnologues sampled from each SSR category (Fig. 3a, b; Supplementary Table 12). However, the three most significantly enriched GOslim (biological process) terms (*cell differentiation*, *signal transduction* and *anatomical structure development*) were shared with ohnologues from SSR1 and/or SSR2 categories. This fits with a broader observation that the majority of overrepresented GOslim terms recorded for ohnologue sets from the SSR categories were shared with the AORe ohnologue set (14 out of 16 for SSR1; 6 out of 8 for SSR2 and 5 out of 8 for SSR345). Terms shared between SSR1 and AORe ohnologues included *growth*, *embryo development* and *cell-cell signalling* (biological process), in addition to *cytoplasm* and *plasma membrane* (cellular components). The terms *vesicle-mediated transport* and *chromosome organization* (biological process) in addition to *nucleolus* and *chromosome* (cellular component) were shared by SSR2 and AORe ohnologues. Terms shared by SSR345 and AORe ohnologues were *cell motility*, *cell adhesion* and *nucleocytoplasmic transport* (biological process) in addition to *enzyme binding* (molecular function). The term *intracellular* (cellular component) was the single term shared between the AORe, SSR1 and SSR2 ohnologue sets. There was little overlap in shared GOslim terms among any of the SSR ohnologue sets, which may reflect a lack of power due to the comparatively smaller number of genes in these sets compared to AORe regions.

Among the GOslim terms unique to the AORe ohnologue set were *cell population proliferation*, *mitotic cell cycle* and *mitotic nuclear division* (biological process), in addition to *lipid binding, enzyme regulator activity*, *kinase activity* and *transcription factor binding* (molecular function). For SSR1 ohnologues, *extracellular matrix organization* (biological process) and *cytoplasmic chromosome* (cellular component) were the only uniquely enriched terms. For SSR2 ohnologues, *plasma membrane organization* was the most significantly enriched term, and was not shared with any other gene set. Unique enriched terms for SSR345 ohnologues were *secondary metabolic process* (biological process) in addition to *peroxisome* and *mitochondrion* (cellular component).

For singletons, only the AORe set showed significant enrichment of GOslim terms, which represented three distinct metabolic processes and two molecular functions, *oxidoreductase activity* and *transmembrane transporter activity*. These terms did not overlap with any of the ohnologue gene sets.

To summarize, these results: i) emphasise differences in gene functional enrichment comparing Ss4R ohnologues and singletons, ii) indicate that shared biases in broadly defined functions (i.e., GOslim terms) between ohnologues with different rediploidization ages/histories are common, and iii) provide evidence for functional enrichment of ohnologues unique to regions with different rediploidization ages, particularly for AORe regions and to a much lesser extent SSR regions.

### Rediploidization age influences gene expression level

A recent study showed that reduced expression level in one or more rarely both ohnologues in a pair was a dominant pathway of evolution following Ss4R (Gillard et al. 2021). This led us to hypothesise that ohnologue expression level will depend on rediploidization age, owing to the amount of evolutionary time ohnologues have had to diverge towards the common outcome of reduced expression. This predicts that the oldest ohnologues in AORe regions will on average show lower expression than younger ohnologues in SSR regions. To test our hypothesis, we compared mean transcript levels quantified by RNA-Seq in a panel of Atlantic salmon tissues (Lien et al. 2016) across the defined rediploidization categories considering both ohnologues and singletons (Supplementary Fig. 14). This revealed that ohnologues indeed often have higher average expression in SSR compared to AORe regions within each tissue (Supplementary Fig. 14). In support of our hypothesis, the ratio of median ohnologue vs. singleton transcript level was significantly reduced in AORe compared to SSR1, SSR2 and SSR345 regions (respective average expression ratio across all tissues: 1.07, 1.51, 1.53 and 2.43) (Fig. 4).

**Fig. 4.**
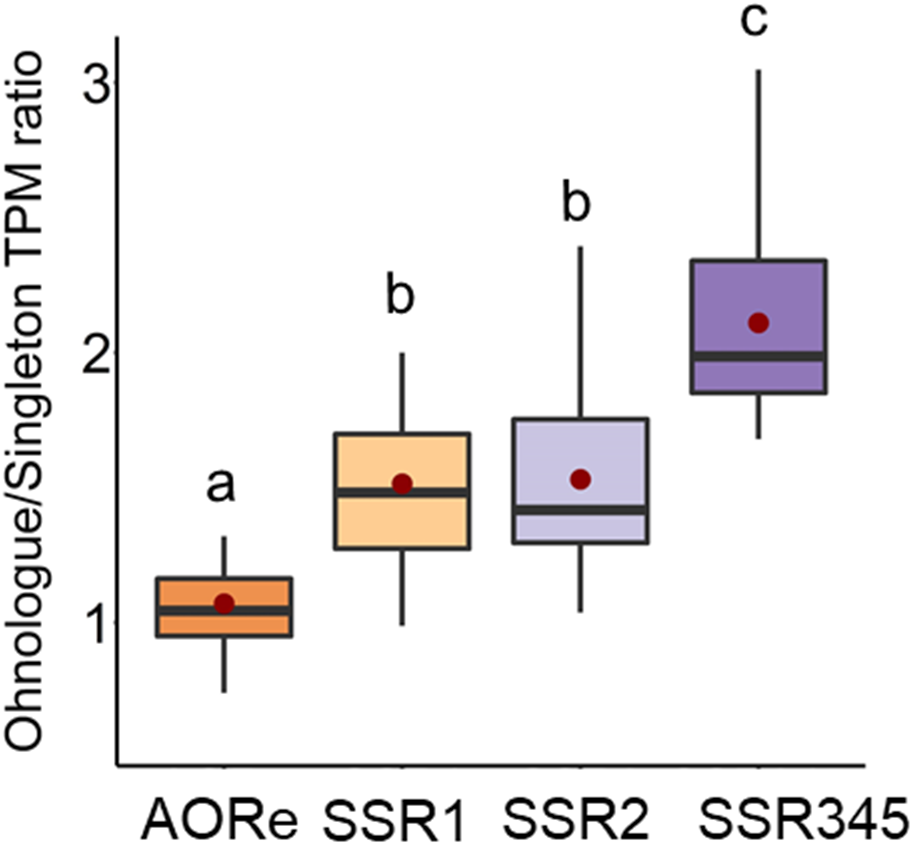
Gene expression level is affected by rediploidization age. The boxplot depicts the ratio of ohnologue to singleton transcript level for genes retained in genomic regions with different rediploidization ages. This summarizes data for fifteen Atlantic salmon tissues, specifically the average per-tissue ratio of ohnologue vs. singleton transcript level (transcripts per million, TPM; from Lien et al. 2016). Boxplot definition: boxes spans the interquartile range, with the median (Q2) as a central bar, the mean as a red circle, and the upper and lower bounds representing the respective minimum and maximum values within the 25th percentile (Q1) and 75th percentile (Q3). Upper and lower whisker bounds are the largest and smallest values within 1.5 times above Q3 and below Q1, respectively. Different letters indicate significant differences according to pairwise T-tests: AORe vs. SSR1: *p* = 0.002; AORe vs. SSR1: *p* = 0.001; AORe vs. SSR1: *p* = 8.3e-8; SSR1 vs. SSR2: *p* = 0.90; SSR2 vs. SSR345 *p* = 0.001.

### Positive selection of ohnologues in regions with distinct rediploidization ages

We previously hypothesised that LORe promotes lineage-specific adaptation by creating a substrate of ‘newly diverging’ ohnologues that can functionally specialize in response to lineage-specific selection pressures (Robertson et al. 2017). We have further argued that selection on ohnologue functions retained in AORe regions may be comparatively constrained by ancestral divergence and specialization inherited prior to speciation (Robertson et al. 2017). To test these ideas, we asked if the number of ohnologues targeted by positive selection was a product of rediploidization age at three distinct periods of salmonid evolution. We employed an established method (Bertolotti et al. 2020; Gillard et al. 2021) to generate codon alignments including Ss4R ohnologues for all species used in our rediploidization analyses, along with additional teleost outgroups to Ss4R. After filtering, we retained 3,355, 709 and 85 alignments in AORe, SSR1 and SSR2 regions, respectively (representing ~8,300 unique ohnologue pairs) (see Supplementary Data 14 for genomic locations; alignments and trees provided in Supplementary Data 7).

Each alignment was used in a *d*N/*d*S analysis employing an adaptive branch-site model (Smith et al. 2015). We documented ohnologues showing evidence for positive selection (corrected *P* < 0.05) comparing each rediploidization age category at three pre-defined phylogenetic branches: [1] ‘post-WGD’, separating the Ss4R event from the divergence of salmonid subfamilies, [2] ‘ancestral Salmoninae’, defining the common ancestor of Salmoninae species and [3] ‘ancestral anadromous Salmoninae’, defining the common ancestor of *Salmo*, *Oncorhynchu*s and *Salvelinus* (Fig. 5a; Supplementary Data 15, 16). This approach was designed to test a prediction of our hypothesis: that more ohnologues will show positive selection in SSR regions (less constrained by ancestral divergence) than AORe regions (more constrained by ancestral divergence) along the tested branches.

**Fig. 5.**
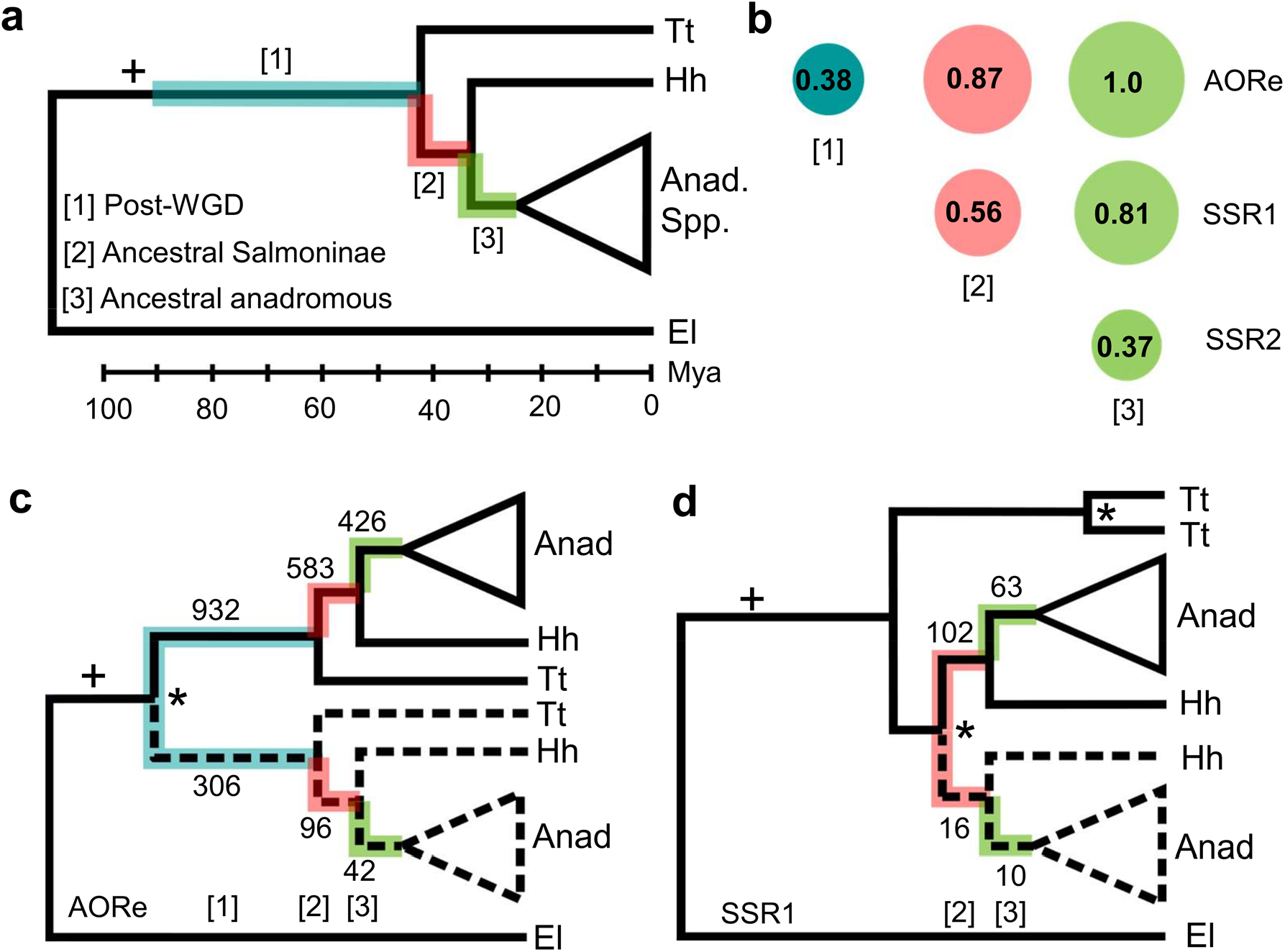
Positive selection on Ss4R ohnologues sampled from regions with distinct rediploidization ages across three periods of salmonid evolution. **a**. Species tree showing lineages used in analysis (Tt = European grayling; Hh = huchen; Anad. Spp = clade consisting of Atlantic salmon, brown trout, rainbow trout, Arctic charr and Coho salmon) highlighting the three test branch categories. **b.** Bubble plots comparing the percentage of ohnologue branches under positive selection (corrected *P* < 0.05) in genomic regions with different rediploidization ages, normalized to millions of years. **c.** and **d.** Ohnologue trees depicting the respective topologies in AORe and SSR1 regions of the genome, highlighted with the number of branches under positive selection. The upper half (solid black line) and lower half (dotted black line) of each tree represents the respective number of branches where a single, or both, ohnologues in each pair was inferred to be under positive selection.

For AORe regions, 16.4%, 12.9%, and 7.9% of the tested ohnologues showed evidence of positive selection along the post-WGD, ancestral Salmoninae and anadromous Salmoninae branches, respectively. To reliably compare across branches, we normalized the effect of absolute time, i.e. the post-WGD branch represents ~43 million years evolution, compared to ~14.8 and 7.9 respective million years for the ancestral and anadromous Salmoninae branches (according to Macqueen and Johnston 2014). This correction indicates that 0.38%, 0.87% and 1.0% of tested ohnologues in AORe regions experienced positive selection per million years along the three respective branches (Fig. 5b). As a caveat of this normalization, we acknowledge that we cannot know when selection occurred along the post-WGD branch, and the presented estimates would be misleading if the majority of positive selection occurred rapidly post-WGD, rather than steadily through time. We observed a similar proportion of ohnologues under positive selection in SSR1 regions for ancestral Salmoninae branches (0.56% per million years) and ancestral anadromous Salmoninae (0.81% per million years) branches. For SSR2 regions, evidence for positive selection was obtained for just five ohnologue genes (i.e. anadromous Salmoninae branches; 0.37% per million years).

To further interrogate how selection targeted ohnologue coding regions at different periods of salmonid evolution, we recorded instances where positive selection affected either one or both of the ohnologues in each pair comparing AORe and SSR1 regions (Fig. 4c, d). In AORe regions, we observed evidence of positive selection acting on both ohnologues in a pair for ~25% of all post-WGD branches, a value significantly higher than ancestral Salmoninae (~14%) or ancestral anadromous Salmoninae (~9%) branches (both *P* < 0.0001; two-sided Fisher’s exact test). For SSR1 regions, positive selection acting on both ohnologues in each pair was inferred for a smaller number of branches than the AORe post-WGD branches in ancestral Salmoninae (~14%) and ancestral anadromous Salmoninae (~14%) branches (*P* = 0.0063 and *P* = 0.034, respectively, two-sided Fisher’s exact test).

## Discussion

This study advances our understanding of the role played by rediploidization dynamics in long-term evolution following autopolyploidization. Our findings, placed alongside the methodological advances reported, have implications for future studies of WGD events. In addition, we have provided useful resources for genomic and evolutionary investigations in salmonids, a group of fishes with extensive ecological and economic importance (Houston and Macqueen 2019; Houston et al. 2020) including a new genome for the endangered huchen (available through the Ensembl browser), genome alignments spanning multiple species inclusive of ohnologue regions, and protein-coding positive selection data for a global set of ohnologue genes.

Delayed rediploidization and LORe are yet to be unequivocally demonstrated outside the salmonid lineage, but were proposed to follow the teleost-specific WGD event (often called Ts3R) and earlier WGD events at the base of vertebrates (Martin and Holland 2014; Robertson et al. 2017; Rozenfeld et al. 2019). There further exists a growing recognition that delayed rediploidization leading to LORe may have followed many of large number of WGD events in plant evolution, most of which are not assigned to autopolyploid or allopolyploid origins (Clark and Donoghue 2017), with several authors proposing that delayed and nested patterns of species diversification are consistent with LORe (Smith et al. 2015; Clark and Donoghue 2017; Van de Peer et al. 2017; Carretero‐Paulet and Van de Peer 2020; Van de Peer et al. 2021). Considering the large and growing number of high-quality genome sequences in diverse eukaryotic lineages, it will be feasible to adapt our genome alignment coupled to phylogenetics approach to address the outcomes of rediploidization for WGD events of a comparable or younger age to Ss4R, exploiting synteny/collinearity to distinguish ohnologues from other gene duplicates. When studying older WGD events, such as Ts3R, genome alignment methods will likely be unsuitable due to the greater evolutionary distances involved. Nonetheless, mapping ohnologue gene tree topologies from multiple species that branched off early following ancestral WGD, to a genome characterised by the same event, should allow for valid tests of LORe. Such an approach can exploit the prior expectation that trees showing the same lineage-specific ohnologue divergence nodes will be co-located within chromosome regions sharing the same ancestral rediploidization histories (Fig. 1) more than expected by chance.

Most evolutionary studies of WGD events use approaches lacking scope to characterize delayed rediploidization. This results from a common assumption that genome-wide ohnologue divergence begins immediately after polyploidization, and will thus be ancestral to any sister lineages sharing the same WGD event. On this basis, many authors have used the distribution of synonymous distance (K_S_) among all paralogous gene pairs within a genome to identify WGD events as peaks against the background distribution(Vanneste et al. 2013). A spectrum of ohnologue divergence ages following delayed rediploidization is not expected to generate a single peak, while distinct rediploidization ‘waves’ (Fig. 2a) would be expected to generate multiple peaks (or tails to peaks), impacting the accuracy of such inferences. Consequently, delayed rediploidization adds to the known limitations of K_S_ methods (Vanneste et al. 2013; Tiley et al. 2018; Zwaenepoel and Van de Peer 2019). Gene tree - species tree reconciliation approaches are widely used to identify WGD events and likewise have recognized caveats (Thomas et al. 2017; Zwaenepoel and Van de Peer 2019). The complex patterns of LORe characteristic of large regions within salmonid genomes (Fig. 1) would strongly violate an assumption common to such methods - that ohnologue divergence starts along a specified branch in a species tree. To further compound this challenge, our data shows that ohnologue trees with the same (i.e., AORe) topology may be associated with very different rediploidization ages (e.g., Ssa09-20 vs. other AORe regions; Fig. 2a). This situation would be invisible to gene tree - species tree reconciliation methods and also has an additional implication for dating WGD events using molecular clocks. When faced with delayed rediploidization, genomic regions where rediploidization occurred earliest provide the best possible estimate for the timing of WGD – because they are ‘as close we can get’ to the WGD event using sequence data (Macqueen and Johnston 2014). Past estimates for the timing of Ss4R either failed to exclude ohnologues showing delayed rediploidization (Berthelot et al. 2014; Lien et al. 2016), or lacked genome-wide information generated in the current study to identify differences in the timing of rediploidization for AORe regions (Macqueen and Johnston 2014). We thus propose that the timing of the Ss4R WGD should be treated as the genomic region with the oldest estimated rediploidization date, i.e. 106 Mya (95% Bayesian credibility interval: 89-125 Mya) (Fig. 2a). While this is only marginally older than a previous reliable estimate based on 18 ohnologue pairs sampled from AORe regions (i.e. 88-103 Mya) (Macqueen and Johnston 2014), the problem would be much larger with greater heterogeneity in the timing of ancestral rediploidization across genomic regions, which may well be the case for other WGD events.

Our study further aimed to expand knowledge on the drivers and outcomes of complex rediploidization histories within a salmonid genome, by combining gene set enrichment, gene expression, and *d*_N_/*d*_S_-based positive selection analyses. We identified a much larger set of enriched GOslim terms for ohnologues than singletons across all tested rediploidization categories and a lack of overlap in functional enrichment across the two gene classes. This emphasises selection on the retention of ohnologues, consistent with past investigations of different WGD events (Blomme et al. 2006; Smet et al. 2013; Inoue et al. 2015; Han et al. 2016; Parey et al. 2020). Previous studies of salmonid genomes have identified major differences in gene functional enrichment comparing ohnologues sampled from AORe vs. SSR regions using standard GO terms (Robertson et al. 2017; Campbell et al. 2019). However, the evolutionary and functional basis for these observations remain unclear. This is partly due to the lack of definition of background expectations, unrelated to rediploidization, comparing gene sets sampled from non-overlapping regions of any genome. In an effort to gain data informative to rediploidization, we used GOslim to capture broadly defined functions represented across non-overlapping genomic regions, along with a stringent cut-off (*P* < 0.001) to capture strong effects. Unlike the previous studies mentioned, we identified extensive overlap in functional enrichment among ohnologues from non-overlapping regions of the Atlantic salmon genome with distinct rediploidization histories. This shared enrichment included several GOslim terms enriched in ohnologues following early WGD events in vertebrates (Makino and McLysaght 2010), linked to development, signal transduction, cell differentiation and cell-to-cell signalling (Fig. 3b). However, we also observed evidence of functional enrichment specific to ohnologues from regions with unique rediploidization histories. This was largely explained by ohnologues from AORe regions, which may partly reflect increased statistical power, as this gene set is markedly larger than ohnologues retained/sampled in SSR regions. However, the result may reflect the greater evolutionary time for selection to act on ohnologue retention, considering that AORe ohnologues have been diverging for tens of millions of years longer than SSR ohnologues (Fig. 2a). Interestingly, many of the enriched GOslim terms specific to AORe regions (Supplementary Data 12) were shared with ohnologues retained from the early WGD events in vertebrates defined as being dosage-balanced, including kinase activity and enzyme regulator activity (Makino and McLysaght 2010). AORe region specific GOslim enrichment may also be linked to selection pressures specific to the early challenges posed by autopolyploidization (Hollister 2015), which may have been resolved in later stages of rediploidization, explaining a lack of enrichment for SSR ohnologues. Perhaps related is our observation that positive selection acted on the coding region of both genes in ohnologue pairs at the highest rate along post-WGD branches in AORe regions. This could reflect a greater need for co-adaptation between protein function and dosage changes in the immediate aftermath of the Ss4R autopolyploidization (Qiao et al. 2019; Gillard et al. 2021), compared to the later stages of rediploidization. Our observation of an inverse scaling in the ratio of ohnologue to singleton expression level with rediploidization age (Fig. 4) is also highly consistent with reports of dosage-related ohnologue downregulation in the post-WGD period of evolution (Gillard et al. 2021). Under this scenario, AORe ohnologues have experienced more evolutionary time to down-tune transcript expression by selection or drift, towards a value closer on average to singleton genes, whereas genes in SSR regions are earlier in the same process.

The small number of GOslim terms enriched specifically in ohnologues from SSR regions are worth highlighting. The lack of enrichment for these terms in the larger AORe ohnologue set is difficult to explain on grounds of lack of statistical power, and our data may thus capture genes functions where rediploidization was selected on during the immediate post-WGD period. Notably for the latest-rediploidization regions (SSR345), where ohnologue sequence divergence is extremely limited (e.g., Fig. 1d, e), we recorded a strong enrichment of two GOslim terms, *mitochondrion* (53 genes; odds ratio: 1.67) and *peroxisome* (11 genes; odds ratio: 3.95). A past study revealed that Ss4R ohnologues showing the most common trajectory of gene expression evolution – where one copy in a pair evolved downregulation along the post-WGD branch - were enriched for equivalent functions using an alternative gene set enrichment tool (Gillard et al. 2021). The mitochondria related enrichment was explained by a possible mismatch in stoichiometric balance between nuclear-encoded ohnologues that interact with mitochondrial-encoded proteins that were not duplicated by WGD. As delayed rediploidization halted ohnologue sequence divergence for at least 50-90 million years in these regions (Fig. 2a), it is plausible that interactions between nuclear and mitochondrial proteins within the energy-generating mitochondria contributed to selection against rediploidization.

Positive selection analysis of ohnologues from regions with different rediploidization histories failed to support our previous hypothesis that LORe boosts adaptation by positive selection on duplicated coding regions. Specifically, we recorded a similar proportion of ohnologues in AORe and SSR1 regions, and limited number of SSR2 ohnologues under inferred positive selection. However, these results may strongly reflect the lack of power to detect positive selection in SSR compared to AORe ohnologue pairs due to their markedly lower sequence divergence (Smith et al. 2015). Irrespective of rediploidization age our analysis highlighted that the rate of ohnologue positive selection was higher during the early period of Salmoninae diversification than for post-WGD branches, with the highest rates noted in the ancestor to anadromous species (Fig. 1a). This warrants follow-up investigations of these ohnologues to understand lineage-specific selection drivers in salmonid evolution, especially with respect to anadromy.

An implication of our work for salmonid genome biology relates to past work that considered all rainbow trout ohnologues classified as belonging to SSR regions in the current study as tetrasomic (Campbell et al. 2019). It is well established that tetrasomic inheritance still occurs in salmonid genomes, particularly impacting males, largely focussed at telomeric regions (Allendorf et al. 2015). Our results warrant caution in confusing delayed rediploidization with an absence of rediploidization. Based on our data, tetrasomic inheritance is unlikely to have occurred for tens of millions of years in regions classified as tetrasomic in rainbow trout (Campbell et al. 2019). These regions have experienced ohnologue divergence descended across various levels of Salmoninae evolution predating the *Oncorhynchus* lineage, including in all SSR1 and SSR2 regions (Fig. 1, 2). We would not expect tetrasomic regions to contain ohnologues showing any sequence divergence. Instead, these regions should harbour up to four alleles, and would collapse in haploid-representative genome assemblies.

A major unresolved question concerns the drivers of delayed rediploidization in salmonid genomes. It could be that rediploidization occurs randomly, but was initially strongly selected against in regions showing delayed rediploidization due to negative impacts for certain genes or due to beneficial impacts of maintaining these genes as tetraploid. Global analyses such as gene set enrichment lack the resolution to resolve such possibilities. Assuming selection acted on rediploidization in SSR regions across tens of millions of years (Fig. 2), our study raises the question of why a second wave of rediploidization was tolerated? As this coincides inextricably with the evolutionary origin of salmonid subfamilies, perhaps the events leading to speciation coincided with reduced effective population size, lowering the efficiency of purifying selection on deleterious impacts of rediploidization. Or perhaps novel selective pressures accompanying early species diversification (e.g. linked to the initial development of anadromy) altered selection on rediploidization and ohnologue divergence through other routes, due to effects on specific genes. The high rate of positive selection on ohnologues at this period of Salmoninae evolution supports this idea indirectly. Mechanistically, rediploidization is linked to a proliferation of TEs in the genome, which cause rearrangements driving the cessation of multivalent meiotic pairings, limiting ohnologue divergence (Soltis et al. 2015). As there have been known bursts of TE activity throughout salmonid evolution (Lien et al. 2016), lineage-specific TE proliferation is likely causatively linked to delayed lineage-specific rediploidization. Unfortunately, the current generation of salmonid genomes do not allow such ideas to be addressed due to their limited representation of TEs and genomic regions showing very recent rediploidization. Emerging long-read assemblies spanning all salmonid genera will allow for more mechanistic insights into the relationship between TE evolution, speciation and lineage-specific rediploidization.

In conclusion, our findings provide a useful model for delayed autopolyploid rediploidization and its macroevolutionary impacts. We advocate for more in-depth investigations of rediploidization dynamics following many other eukaryotic WGD events, potentially demanding the uptake or creation of phylogenomic methods that better accommodate the expectations of delayed and lineage-specific rediploidization. Such work will be essential to define the prevalence and significance of LORe and delayed rediploidization in wider evolution.

## Materials and Methods

### Huchen genome assembly and RNA sequencing

Full methods are provided in Supplementary Methods 1. Briefly, sampling was done using genetically wild hatchery reared fish from the State Fisheries Department Farm, Lindbergmühle, Germany. The embryo used for sequencing was generated from wild parents, with haploidy induced by UVC irradiation. Genomic DNA was extracted from a confirmed haploid (Supplementary Fig. 1) and used to construct paired-end (500bp insert) and mate-pair sequencing (~6kb and ~12kb insert libraries) (Heavens et al. 2015) libraries. Sequencing was done using an Illumina HiSeq2500 with 250 bp paired-end reads. Genome size was estimated using a k-mer approach (Vurture et al. 2017). Contig assembly, scaffolding and gap-filling were done using W2RAP-CS42_TGACv1 (Clavijo et al. 2017), SSPACE v3.0 (Boetzer et al. 2011) and GapFiller 1.10 (Boetzer et al. 2011), respectively. Assembly completeness and quality was estimated using CEGMA v2.5 (Parra et al. 2007), BUSCO v.3 (Waterhouse et al. 2018) and KAT tools v.2.3.4 (Mapleson et al. 2017). Repeat modelling and masking was performed using Repeatmodeler v1.0.9 (Smit and Hubley 2015) and Repeatmasker v4.0.7(Smit et al. 2015) (Supplementary Table 3).

218Gb RNA-Seq data (~720 million paired-end 150bp reads) was generated to support annotation of the huchen genome (available through the NCBI BioProject PRJNA480959). The samples represented fifteen tissues from one individual (fork-length: 30cm, sex not identifiable), namely whole eye, whole mixed brain, swim bladder, gill filament, olfactory pit, skin, skeletal muscle, stomach, distal intestine, unidentified gonad, pyloric caeca, kidney, spleen, liver, and heart). We also generated liver and unidentified gonad data for a further three individuals (fork-length: 28-31cm; sex not identifiable). Total RNA was extracted using Trizol (Sigma) following the manufacturer’s protocol. Library construction was carried out using an Illumina TruSeq RNA kit and sequencing performed on a HiSeq1500 platform by the Norwegian sequencing centre. An annotated version of the huchen genome with 50,114 protein coding gene predictions is available on the Ensembl genome browser (https://www.ensembl.org/Hucho_hucho).

### Whole genome alignment capturing ohnologues

We developed an approach to circumvent the fact that genome alignment tools are geared towards orthologous regions and not designed to capture multispecies ohnologue variation (summarized in Supplementary Fig. 15). This approach leverages prior knowledge of collinear/syntenic ohnologue blocks retained from WGD and inputs ohnologue sequence variation to the alignment algorithm as different ‘species’, allowing multispecies alignments to be generated inclusive of ohnologues. For AORe regions, genome assemblies for Atlantic salmon (Lien et al. 2016), European grayling (Varadharajan et al. 2018), huchen (this study), Chinook salmon (*O. tshawytscha*) (Christensen, Leong, et al. 2018), rainbow trout (Pearse et al. 2019) and northern pike (Rondeau et al. 2014) were used. For SSR regions, we added a *Salvelinus* genome (NCBI accession: GCA_002910315.2) to increase scope to capture LORe outcomes.

The first step was to identify sequences homologous to the two established ohnologue blocks in Atlantic salmon (Lien et al. 2016) (Supplementary Data 1 and 2 for AORe and SSR regions, respectively) separately for each target species. While chromosome anchored sequence was used for the Atlantic salmon reference, we used scaffolds for other species to recover maximal data, either because no chromosome level assembly was available (e.g. huchen and European grayling), or a large number of scaffolds were not anchored to chromosome. We generated BLASTn (Altschul et al. 1997) databases (using the makeblastdb module) for each defined ohnologue block in the Atlantic salmon genome (Lien et al. 2016) and performed per species BLASTn searches using genome scaffolds as queries (e-value cut-off: 0.001, maximum target sequences: 3, max_hsps = 20000, word size of 40 and minimum 90% sequence identity) before filtering the hits for minimum alignment length (<3,000 bp) and linearity using a published script (Christensen, Rondeau, et al. 2018). This step captures high quality scaffolds sharing close homology to the two Atlantic salmon reference ohnologue sequences for each species, which were retrieved as fasta files using fasta_tools within MAKER v3.0 (Cantarel et al. 2008).

The next step involved splitting the recovered scaffolds in each species homologue set into two files representing the distinct ohnologue sequences. This is crucial to allow genome alignment tools to accept two ohnologues per species within the same alignment. For AORe regions, we performed an initial step to categorise scaffolds on the basis of putative 1:1 orthology to each Atlantic salmon ohnologue, which is possible due to ancestral rediploidization (Macqueen and Johnston 2014; Robertson et al. 2017). This was done by aligning all retrieved scaffolds per ohnologue block to a reference containing both Atlantic salmon ohnologue sequences using Mugsy v1.2.3(Angiuoli and Salzberg 2011). We retrieved the number of alignment locations per scaffold to each Atlantic salmon ohnologue from the resultant MAF file using a custom script (Supplementary Methods 2). This allows us to identify the likely orthologous scaffold to each Atlantic salmon ohnologue under the rationale that orthologous sequences share more alignment locations. In most cases, all scaffold alignment locations matched to a single Atlantic salmon ohnologue as expected. We excluded any (possibly chimeric) scaffolds where <70% of the alignment locations matched to a single Atlantic salmon ohnologue. At the end of this step, we split each set of scaffolds per species into two fasta files representing two ohnologue sets and renamed the fasta headers to represent orthology with one of the two Atlantic salmon ohnologue regions (i.e. ‘species abbreviation_Atlantic salmon chromosome arm name_scaffold name’).

For SSR regions, it is either challenging (due to recent ancestral rediploidization) or impossible (due to LORe) to identify 1:1 orthology for ohnologue sequences across salmonid species (Robertson et al. 2017). Consequently, we modified our approach to bin scaffolds homologous to the Atlantic salmon ohnologue blocks (Supplementary Data 2) into two groups that individually contain no more than one ohnologue per species, allowing them each to be included in the genome alignment. To achieve this, all scaffolds homologous to each Atlantic salmon ohnologue pair were aligned to one of the Atlantic salmon sequences using Minimap2 v2.16(Li 2018) (kmer size: 27; ≤5% sequence divergence allowed between subject and reference; other parameters default). Alignments were visualized in Tablet v.1.17.08.17(Milne et al. 2010) allowing scaffolds to be binned into two groups of ohnologue per species based on shared overlap of two distinct ohnologues to a single Atlantic salmon reference. Where a single alignment was present, potentially due to assembly collapse (Varadharajan et al. 2018), scaffolds were randomly binned into one of the two ohnologue groups. We renamed the ohnologous scaffolds in the two bins as for AORe regions, except using either ‘Chr-A’ or ‘Chr-B’ designations to replace Atlantic salmon chromosome arm names.

The final step was to align all sequences homologous to each ohnologue block in Atlantic salmon across the different species. This was done using Mugsy v1.2.3 (Angiuoli and Salzberg 2011), setting one of the Atlantic salmon ohnologue sequences as the reference, and then adding separate fasta files to the alignment for i) the other Atlantic salmon ohnologue sequence, ii) two distinct sets of ohnologous scaffolds per salmonid species, and iii) a single set of co-orthologous scaffolds for northern pike. The maximum distance along a single sequence to chain the anchors into a single local collinear block (LCB) was set to 2,000 bp, and the minimum span of aligned regions within LCBs was set to 100bp (other parameters default). Summary statistics for the final MAF files produced for AORe and SSR regions (Supplementary Data 3 and 4, respectively) were extracted using mafStats within mafTools v01(Earl et al. 2014).

### Alignment processing and phylogenetics

We processed the alignment blocks within each MAF file to capture maximal useful information on ohnologue evolution for phylogenetic analysis. This was done by filtering on the basis of the number of represented sequences (a product of the different taxa plus Ss4R ohnologues captured) using a custom script (Supplementary Methods 2). For AORe regions, alignments were filtered for a minimum of nine out of eleven possible sequences in any block, meaning most salmonid species were required to retain two ohnologues, leading to a total of 92.2 Mb alignment blocks of which 34,603, 43,329 and 28,613 included eleven, ten and nine sequences, respectively (Supplementary Data 5). For SSR regions, we allowed a more inclusive filtering strategy to capture data in regions where rediploidization was most delayed (i.e. ohnologues have diverged the least) and assembly collapse (Varadharajan et al. 2018) and fragmentation is common. In SSR regions, up to thirteen sequences were possible across different taxa and retained ohnologues; we recovered 15,313 alignments with >11 sequences represented, and a further 7,805 alignments with >9 sequences represented (Supplementary Data 6).

The parsed MAF files were converted to fasta format using Maffilter v1.3.1 (Dutheil et al. 2014). The fasta-splitter script (Lam et al. 2015) was then used to split each alignment block per MAF file into individual fasta files. Each of the split fasta files was processed through GBlocks v0.91b (Castresana 2000) using default parameters to filter low quality alignment regions. These finished alignments were used to construct maximum likelihood phylogenetic trees in IQTREE v1.6.8(Nguyen et al. 2015), using the best fitting substitution model (Kalyaanamoorthy et al. 2017) and ultrafast bootstrapping (Minh et al. 2013) with 1,000 iterations to obtain branch support values.

### Reconstructing rediploidization history and LORe in SSR regions

We matched expectations of LORe (Robertson et al. 2017) against empirical data on ohnologue divergence captured by phylogenetic trees sampled from SSR regions. All trees were assigned to one of the five possible SSR categories (SSR1-5; Fig. 1b), facilitated using scripts executed in R (R core team, 2020) Supplementary Methods 2) followed by manual checking of every tree. All trees were rooted to northern pike, or European grayling when pike was absent from the alignment, or in rare situations where both species were absent, using midpoint rooting. To initially assign trees into different SSR categories, we exploited predicted monophyly for included species (Fig. 1b), along with the *Dupfinder* (Varadharajan et al. 2018) function to confirm the position of nested ohnologue clades. This allowed us, for example, to initially assign trees to SSR1 on the criteria of i) monophyly of European grayling and ii) a duplication node shared by all Salmoninae members; or to SSR2 on the criteria of i) separate monophyly of both European grayling and huchen and ii) a duplication node shared by all ancestrally anadromous Salmoninae members (and so on for SSR3, 4 and 5). After binning the trees into the five categories, all trees were plotted and manually visualized to check the automatic assignments. At this point, we removed ambiguous trees on the basis of unexpected branching and/or bootstrap values <50 along inferred rediploidization nodes. In some cases, we observed trees where a species showed monophyletic ohnologue sequences with strong bootstrap support, yet clustered within one of two ohnologue clades present elsewhere in the tree. This occurred commonly for trees assigned to SSR2 (i.e., two monophyletic huchen sequences branched as a sister to one of two ohnologue clades representing the ancestrally anadromous Salmoninae species). We accepted such topologies as consistent with SSR2. We present examples of accepted trees for each SSR category in Supplementary Fig. 16-20, while all 11,139 trees used in Fig. 1 are provided in Supplementary Data 7.

A custom script (Supplementary Methods 2) was used to retrieve Atlantic salmon chromosome coordinates for all SSR trees (Supplementary Data 8). The data was visualised as circos plots using OmicCircos v1.26.0 (Hu et al. 2014) or Circlize v0.4.11 (Gu et al. 2014). Supplementary Methods 3 describes the positional mapping of phylogenetic trees with different SSR topologies against brown trout chromosomes homologous to Atlantic salmon ohnologue blocks 03-06.

### Estimating rediploidization age across the genome

To estimate rediploidization age across defined ohnologue blocks within the Atlantic salmon genome (Lien et al. 2016), we used a concatenation approach to maximise the available data. For AORe regions, all alignments per defined ohnologue block were concatenated using SeqKit v0.8.0 (Shen et al. 2016) to generate a single alignment file (described in Supplementary Data 10). For SSR regions, we concatenated alignments across defined ohnologue blocks to capture each inferred SSR category (Fig. 1b; Supplementary Fig. 2-10), i.e. separate concatenations were generated for regions inferred as SSR1, 2, 3, 4 and 5 (Supplementary Data 10). Each sequence file was aligned using Mafft v7.0 (Katoh and Standley 2016) with default parameters. MCMCtree (within PAML-v4.9h) (Yang 2007) was used to estimate rediploidization age, represented by the divergence time of ohnologue sequences (Macqueen and Johnston 2014), using approximate likelihood, an independent rate clock model (clock=2) and the general time reversable substitution model (model=7), allowing independent rates for all nucleotide substitutions. An input tree topology was used to set the appropriate rediploidization history (e.g. AORe, SSR1 etc.) and temporally constrained using uniform distribution priors (after Macqueen and Johnston 2014) (visualized in Supplementary Fig. 21). Each analysis was allowed to run for 30 million generations with a burn-in value of 100,000/30,000 and sample frequency of 1,000/20 for AORe and SSR alignments, respectively (leading to respective sample sizes of 30,000 and 100,000). The mean Bayesian divergence times and 95% credibility intervals estimated by MCMCtree for each rediploidization node were plotted using ggplot2 v3.3.2(Wickham et al. 2016).

### Gene set enrichment analyses

We used an R script (Supplementary Methods 2) to: i) extract all annotated genes in the Atlantic salmon ICSASG_v2 genome GFF file according to coordinates defining regions with distinct rediploidization ages - generating sets of genes combined across different AORe, SSR1 and SSR2 regions, and combining across all SSR3, 4 and 5 regions (respective coordinates for AORe and SSR regions given in Supplementary Data 1 and Data 10) and ii) cross-reference each gene set with the high-confidence Ss4R ohnologues and singleton genes defined in Bertolotti et al. 2020, removing any non-matching genes. GO enrichment tests on these gene sets were done (pipeline described in https://gitlab.com/cigene/R/Ssa.RefSeq.db/-/wikis/go-slim), against all Atlantic salmon genes as the background. Enriched GOslim terms from each gene set were parsed to generate a list of unique and overlapping GOslim terms using an R script (Supplementary Methods 2), and the results visualised through a network graph generated in ggnetwork v.0.5.8 (Briatte 2020).

### Gene expression analysis

RNA-Seq derived gene expression values (transcripts per million, TPM) across fifteen tissues (Lien et al. 2016) for the defined sets of ohnologues and singletons were retrieved using *salmonfisher* (https://gitlab.com/sandve-lab/salmonfisher) and plotted using ggplot2 v3.3.2(Wickham et al. 2016). The median TPM values per tissue were used to calculate ohnologue vs singleton TPM ratios (Fig. 4), which were compared across gene sets using paired t-tests.

### Positive selection analyses using Ss4R ohnologues

For the positive selection analyses of ohnologues from AORe, SSR1 and SSR2 regions, curated protein-coding alignments and trees were generated using a published pipeline (Bertolotti et al. 2020) (scripts available at https://gitlab.com/sandve-lab/salmonid_synteny). Briefly, Orthofinder v.2.4.0 (Emms and Kelly 2019) was used to generate orthogroups inclusive of Atlantic salmon, brown trout, rainbow trout, coho salmon, Arctic char, huchen, European grayling, northern pike, three spined stickleback, medaka and Nile tilapia. Coding sequences were extracted from each orthogroup and aligned using Macse v.2.03(Ranwez et al. 2011) before gene trees were generated using TreeBeST v.1.9.2 (https://github.com/Ensembl/treebest). The alignments were split into subtrees according to the presence of pike as an outgroup to salmonid clades. Trees falling within AORe regions (Supplementary Data 1) were filtered by additional criteria: i) the presence of the ancestral Ss4R node with bootstrap support >70 and ii) the branching of one or both the European grayling and huchen salmon ohnologues in the correct position with bootstrap support >70. Trees representing SSR regions (Robertson et al. 2017) (Supplementary Data 2) were manually categorised as SSR1 or SSR2. The codon alignments were used to generate evidence of positive selection using the adaptive Branch-Site Random Effects Likelihood model (Smith et al. 2015) within command line HyPhy v2.5.9 (Kosakovsky Pond et al. 2005). Branch-specific *d*N/*d*S values and *P*-values indicative of significant positive selection (corrected *p*<0.05) derived from a likelihood ratio test (Smith et al. 2015) were retrieved in tabular format for branches of interest (i.e. data in Fig. 4) using a custom R script (Supplementary Methods 2).

## Author contributions

MKG and DJM designed the research with inputs from SRS, DH, SAM and SL. JG coordinated the huchen sampling and dissected fish. MKG sampled huchen embryos and led the huchen genome assembly. MKG developed the genome alignment approach and performed phylogenomic analyses with help from TTH and LG. MKG and DJM designed figures, tables and the supplementary information. MKG and DJM co-wrote the manuscript with input from all authors leading to the submitted manuscript.

## Acknowledgements

This work was supported by the Biotechnology and Biological Sciences Research Council grant BBS/E/D/10002070 and the Frimedbio program of the Research Council of Norway (grant number 241016). MKG received studentship funding from a University of Aberdeen Elphinstone scholarship with additional support from the Government of Karnataka. We thank Dr Sebastian Beggel, Dr Bernhard C. Stoeckle, Jens-Eike Täuber and Ms Haiyu Ding at the Aquatic Systems Biology Unit, Technical University of Munich for their support in sampling huchen. We thank Dr Torfinn Nome for supporting bioinformatic analyses. The Earlham Institute performed library preparation and sequencing used in the huchen genome assembly.

## Data availability

All data supporting the findings of this study are available within the paper and its supplementary files, including supplementary data provided through Figshare at https://figshare.com/s/b30a7c7a579392320085. The huchen genome assembly is available through NCBI (accession number: GCA_003317085.1; https://www.ncbi.nlm.nih.gov/assembly/GCA_003317085.1/) and the Ensembl Genome Browser (https://www.ensembl.org/Hucho_hucho). RNA-Seq data produced for huchen samples is available through NCBI (BioProject: PRJNA480959; https://www.ncbi.nlm.nih.gov/bioproject/PRJNA480959). All scripts used in the study are provided in Supplementary Methods 2 or are otherwise accessible through links provided in the Methods section.

