## Supplementary information for "Genome-wide reconstruction of rediploidization following autopolyploidization across one hundred million years of salmonid evolution"

Contents:

**Supplementary Methods**

Supplementary Methods 1: Generation of a draft huchen genome assembly
Supplementary Methods 2: Scripts used in the study
Supplementary Methods 3: Mapping SSR trees to the brown trout genome

**Supplementary Figures:**

Supplementary Fig. 1. Huchen genome assembly validation
Supplementary Fig. 2. Mapping of reconstructed lineage-specific rediploidization histories for ohnologue regions Ssa03 and Ssa06 in the Atlantic salmon genome.
Supplementary Fig. 3. Mapping of reconstructed lineage-specific rediploidization histories for ohnologue regions Ssa04 and Ssa08 in the Atlantic salmon genome.
Supplementary Fig. 4. Mapping of reconstructed lineage-specific rediploidization histories for ohnologue regions Ssa02 and Ssa12 in the Atlantic salmon genome.
Supplementary Fig. 5. Mapping of reconstructed lineage-specific rediploidization histories for ohnologue regions Ssa02 and Ssa05 in the Atlantic salmon genome.
Supplementary Fig. 6. Mapping of reconstructed lineage-specific rediploidization histories for ohnologue regions Ssa03 and Ssa23 in the Atlantic salmon genome.
Supplementary Fig. 7. Mapping of reconstructed lineage-specific rediploidization histories for ohnologue regions Ssa07 and Ssa17 in the Atlantic salmon genome.
Supplementary Fig. 8. Mapping of reconstructed lineage-specific rediploidization histories for ohnologue regions Ssa11 and Ssa26 in the Atlantic salmon genome.
Supplementary Fig. 9. Mapping of reconstructed lineage-specific rediploidization histories for ohnologue regions Ssa16 and Ssa17 in the Atlantic salmon genome.
Supplementary Fig. 10. Mapping of reconstructed lineage-specific rediploidization histories for ohnologue regions Ssa05 and Ssa09 in the Atlantic salmon genome.

Supplementary Fig. 11. Mapping of reconstructed lineage-specific rediploidization histories for ohnologue regions Str01 and Str32 in the brown trout genome (orthologous to Ssa06 and Ssa03 in the Atlantic salmon genome, respectively).
Supplementary Fig. 12. Mapping of reconstructed lineage-specific rediploidization histories contrasting Ssa06 from Atlantic salmon with its orthologous region in brown trout (Str01)
Supplementary Fig. 13. Mapping of reconstructed lineage-specific rediploidization histories for ohnologue regions Ssa03 and Ssa06 in the Atlantic salmon genome, using protein-coding gene trees as opposed to our genome-wide alignments.
Supplementary Fig. 14. Gene expression of ohnologue and singleton gene sets residing in genomic regions with distinct rediploidization ages
Supplementary Fig. 15. Summary of steps used to generate multispecies ohnologue alignments in advance of phylogenomic analyses.
Supplementary Fig. 16. Example trees for SSR1 topology
Supplementary Fig. 17. Example trees for SSR2 topology
Supplementary Fig. 18. Example trees for SSR3 topology
Supplementary Fig. 19. Example trees for SSR4 topology
Supplementary Fig. 20. Example trees for SSR5 topology
Supplementary Fig. 21. Temporal constraints used in Bayesian rediploidization age analysis for genomic regions with distinct rediploidization histories

**Supplementary Tables**

Supplementary Table 1. Statistics for huchen genome sequencing data
Supplementary Table 2. Statistics for the final huchen genome assembly
Supplementary Table 3. Repeat prediction within the huchen genome

**Supplementary Data -** Provided at https://figshare.com/s/b30a7c7a579392320085

Supplementary Data 1: Position of ohnologue blocks in ancestral rediploidization (AORe) regions within the Atlantic salmon genome
Supplementary Data 2: Position of ohnologue blocks in subfamily-specific rediploidization (SSR) regions within the Atlantic salmon genome
Supplementary Data 3: Multiple alignment format file statistics for ohnologue blocks within ancestral rediploidization (AORe) regions (prior to data filtering)
Supplementary Data 4: Multiple alignment format file statistics for ohnologue blocks within subfamily-specific rediploidization (SSR) regions (pre data filtering)

Supplementary Data 5: Number of alignments retained within ancestral rediploidization (AORe) regions post filtering
Supplementary Data 6: Number of alignments retained within subfamily-specific rediploidization regions post filtering
Supplementary Data 7: All sequence alignments and phylogenetic trees used in the study
Supplementary Data 8: Information on sequence alignments and phylogenetic trees used in rediploidization analyses for AORe and SSR regions (accompanying alignments and tree files are provided in Supplementary Data 7).
Supplementary Data 9: Ohnologue gene tree topologies mapped to different predicted subfamily-specific rediploidization (SSR) scenarios.
Supplementary Data 10: Details of concatenated alignments used in MCMCtree Bayesian analysis to date the onset of ohnologue divergence (i.e. rediploidization ages) across AORe and SSR regions defined in the Atlantic salmon genome.
Supplementary Data 11: Estimated Bayesian divergence times for the onset of ohnologue divergence (i.e. rediploidization ages) across the concatenated alignments (Supplementary Data 9) for duplicated regions in AORe and SSR regions.
Supplementary Data 12: Functional enrichment in biological processes (GOslim terms) for ohnologue pairs and singleton genes extracted from regions of the genome with unique rediploidization histories.
Supplementary Data 13: Genes used in GOSlim analyses detailed in Supplementary Data 12 (matched to the same categories described therein)
Supplementary Data 14: Details of alignments used in ohnologue positive selection analyses
Supplementary Data 15: Results of Branch-Site Random Effects Likelihood test for ohnologue selection across three different rediploidization categories contrasting three defined branches of salmonid phylogeny
Supplementary Data 16: Ohnologue genes under positive selection along three defined branches of salmonid phylogeny according to Branch-Site Random Effects Likelihood test for ohnologues across three different rediploidization categories (i.e. Supplementary Data 15)

#### Supplementary Methods

##### Supplementary Methods 1: Generation of a draft huchen genome assembly

*Haploidy induction*

Haploidy was induced using a bespoke UVC chamber. High-quality eggs from four females and milt from three males was collected into sterile falcon tubes. The milt was chilled on ice, pooled and diluted with an extender (7.25 g NaCl, 0.4 g KCl, 0.8 g NaHCO3, 2 g glucose mixed in 1L distilled water) (1:4 Milt:Extender)(Şahin et al. 2013). The diluted milt showed high motility under a light microscope. The

UVC bulb (254nm) was allowed to reach maximum intensity before use, and a UVC meter was used to measure the intensity of irradiation. Diluted spermatozoa were spread on petri dishes forming thin layers of ~1mm. Separate replicated petri dishes were irradiated for 3, 6, 9 and 12 mins at an intensity of 1464 uw/cm2, with a distance of 10cm between the spermatozoa and UVC bulb. Irradiated spermatozoa were used to fertilize four different batches of eggs. The fertilized eggs were incubated at 11ºC under a flow-through system and monitored daily to remove any dead eggs. UVC treated eggs were sampled 23d post-fertilization when the eggs in a control groups without UVC irradiation started to hatch. Eggs were fixed in 95% ethanol and stored at -20ºC. Embryos subjected to irradiation from all time points, along with control diploids, were analysed for relative DNA content by flow cytometry using eye cells (done by Xelect Ltd, St Andrews) (Fig. S1). The UVC treated embryos were under-developed and a major proportion remained unhatched. Flow cytometry analysis confirmed haploidy in embryos subjected to irradiation for 3 and 6 mins.

*Genomic DNA extraction and sequencing*

Genomic DNA (gDNA) was extracted from candidate haploid embryos using the Thermo Scientific® Genomic DNA Purification Kit (K0512) and eluted in ~100µl of 1X TE buffer. Concentration was measured using a Qubit® 2.0 fluorimeter (dsDNA HS ASsay Kit, Invitrogen™), integrity was assessed using agarose gel electrophoresis, and purity was assessed using a Nanodrop 2000 system (Thermo Scientific®). The sequenced haploid gDNA sample exhibited 260/230 and 260/280 nm ratios of 1.9 and 1.9 and had a total yield and concentration of 6.55 ug. The sample displayed high integrity on an agarose gel. The gDNA sample was sent to the Earlham Institute (Norwich, UK), where paired-end and mate-pair libraries were constructed. Prior to library creation, the gDNA was run on a TapeStation 4200 system (Agilent Technologies) to confirm its integrity and its concentration was re-quantified as described above. A PCR-free library (~500bp insert) was prepared using the TruSeq DNA Sample Prep kit (Illumina) and sequenced on 2.5 lanes of the Illumina HiSeq 2500 with a 250bp paired-end metric. In addition, two mate-pair libraries of ~6kb and ~12kb were generated using the method outlined in Heavens et al. 2015 and sequenced on one HiSeq 2500 lane, again with a 250bp paired-end metric. Full statistics on the raw sequence data are provided in Supplementary Table 1.

*Quality control and trimming*

Sequencing data quality control was done using FastQC (version 0.11.3) (Andrews 2010). Paired-end reads were quality trimmed using TrimGalore (https://github.com/FelixKrueger/TrimGalore) so that all bases had a quality score >30, removing any read (and its pair) trimmed to less than 40 bp length. Adapter trimming was performed, setting the --stringency parameter to 5. Mate pair reads were trimmed using the same approach and subjected to further mate-pair adapter trimming using NxTrim (O'Connell et al., 2015) set to retain sequences ≥40bp post-trimming. This ensured high-quality mate-pair reads were separated from those devoid of mate-pair adapters or showing the wrong orientation. Statistics for quality-controlled final sequence data are presented in Supplementary Table 2.

*Genome size estimation*

Genome size was estimated using the paired end sequence data. Jellyfish-2.2.6 (Marçais and Kingsford 2011) was used to generate k-mer distribution histograms for k-mer sizes of 101 and 111 using ~100M QC-passed paired reads. KAT tools v.2.3.4 (Mapleson et al. 2017) was used to generate a k-mer distribution histogram for a k-mer size of 67 with the 'KAT comp' option. The k-mer distributions were analysed via Genomescope (Vurture et al. 2017), which predicted a genome size of 2.0 to 2.2 Gb for the Jellyfish histograms and 1.9–2.1Gb for the KAT histogram. A single homozygous peak was observed in the k-mer distribution histograms generated through both approaches, providing additional evidence for haploidy (Supplementary Fig 1).

*Genome assembly*

Genome assembly was performed in a stepwise process using different algorithms for contig assembly, scaffolding and gap-filling. Contig generation done using W2RAP (Clavijo et al. 2017) with a k-mer value of 208, minimum contig length filter of 800bp and remaining parameters set to default. The contig assembly was evaluated for completeness using BUSCO v3 against the Actinopterygii dataset (Simão et al. 2015) and contiguity statistics were generated using the abyss-fac option within the ABySS 2.0 toolkit (Jackman et al. 2017). This initial assembly had a size of 1.83 Gb comprising 268,622 contigs and a BUSCO (Waterhouse et al. 2018) completeness score of 74.9% (see next section). Scaffolding was done using the mate-pair sequence data with SSPACE 3.0 (Boetzer et al. 2011). The W2RAP contigs were scaffolded in succession using the 6Kb followed by the 12Kb insert libraries. The assembly was then gap-filled using GapFiller 1.10 (Boetzer and Pirovano 2012) for four iterations using 80x coverage paired-end sequence data. Scaffolding using SSPACE 3.0 resulted in a marked increase in assembly size and >14-fold increase in contiguity (Supplementary Table 2). Gap closing with four iterations eliminated a total of 66.43 Mb spanning 34,132 gaps. The overall gap content in the assembly remained at around 569 Mbp. Statistics for the final assembly are given in Supplementary Table 2.

*Genome completeness assessment*

The scaffolded huchen assembly was screened for 458 conserved eukaryotic proteins using CEGMA (Parra et al. 2007) and 4,584 conserved Actinopterygii genes using BUSCO (Waterhouse et al. 2018). The assembly achieved a CEGMA completeness score of 95% (Supplementary Table 2). 4,135 (90.2%) of the BUSCO genes were complete in the final assembly (Supplementary Table 2). Fragmented BUSCOs comprised a small (3.2%) proportion of genes in the assembly. Assembly quality was further assessed using KAT tools v.2.3.4.(Mapleson et al. 2017). KAT comp was used to compare k-mer content of the paired-end reads to the k-mer content of the scaffolded assembly. A spectra plot was used to gauge completeness and the presence of mis-assemblies, which displayed a normal distribution, consistent with a haploid assembly (Supplementary Table 2).

*Repeat prediction and masking*

Repeat masking was performed using RepeatMasker (Smit et al. 2015). Repbase (Bao et al. 2015) was used to download updated repeat databases (44,968 sequences, 117.2 Mb). Repeat libraries for Atlantic salmon (http://lucy.ceh.uvic.ca/repeatmasker/cbr_repeatmasker.py) and European grayling (Varadharajan et al. 2018) containing 2,494 (1.2 Mb) and 1,195 (0.87 Mb) respective sequences were also downloaded. In addition, de novo repeat modelling was performed using RepeatModeler (Smit et al. 2015; Smit and Hubley 2015). The de novo repeat library was subjected to blastx (Altschul et al. 1997) against the UniProt database to remove any repeats matching an annotated protein-coding gene. All these datasets were merged into a single library that was used for the repeat masking. The curated de novo repeat library salmon contained 1,158 sequences comprising 0.9 Mb. Repeat masking in combination with the other repeated libraries masked a total of 932.1 Mb (~37 %) of the huchen genome. Unclassified elements dominated the different repeat categories accounting for ~ 429 Mb (Supplementary Table 3).

**Supplementary Methods 2: Scripts used in study**

```
####################################################################
script: Filter a MAF file for alignment depth of two and retrieve count data for each alignment.##(BASH)
##################################################################
#1
for i in *.maf
do
grep -A 2 "mult=2" $i >$i.filter
done #pulls out all the alignment blocks with depth=2 from a maf file
#2
for i in *.filter
do
grep -A 1 -B 1 "ChrA" $i >$i.ChrA _mult2;
done #split the maf file into separate files for each Atlantic salmon ohnologue sequence.
#3
for i in *mult2
do
grep ‘scaffold|NW’ $i | \ #
cut -c 1-50 | \
cut -f 1 | \
cut -f 2 -d '.' | sort | \
uniq -c >$i.ids #generate counts for each scaffold in each MAF file.
done

##############################################################################
Categorise trees into different SSR levels (R script)
```

```
##########################################################
library(ape)
library(phytools)
library(phangorn)
library(geiger)

check.topologya = function(tree){
  salmonid.tips = c('huc', 'ssa', 'omy', 'oki', 'ots','thy', '1sa', 'sal')
  if(length(grep('thy', tree$tip.label, value = T))==0) cat ("no_grayling\n")
  if(is.monophyletic(tree,grep('thy',tree$tip.label[substr(tree$tip.label, 1, 3) %in% salmonid.tips],invert = T,
value = T))){
    cat('LORe_thy\n')}
  treex<-drop.tip(tree, grep('thy', tree$tip.label, value = T))
  if(length(grep('huc', treex$tip.label, value = T))==0) cat ("no_hucho\n")
  if(is.monophyletic(treex,grep('huc',treex$tip.label[substr(treex$tip.label, 1, 3) %in% salmonid.tips],invert
= T, value = T))){
    cat('LORe_huc\n')}
  treey<-drop.tip(treex, grep('huc', treex$tip.label, value = T))
  if(length(grep('ssa', treey$tip.label, value = T))==0) cat ("no_salmo\n")
  if(is.monophyletic(treey,grep('ssa',treey$tip.label[substr(treey$tip.label, 1, 3) %in% salmonid.tips],invert
= T, value = T))){
    cat('LORe_ssa\n')}
  treez<-drop.tip(treey, grep('ssa', treey$tip.label, value = T))
  if(length(grep('sal', treez$tip.label, value = T))==0) cat ("no_salvelinus\n")
  if(is.monophyletic(treez,grep('sal',treez$tip.label[substr(treez$tip.label, 1, 3) %in% salmonid.tips],invert =
T, value = T))){
    cat('LORe_sal\n')
     tree = drop.tip(tree, grep('huc', tree$tip.label, value = T))
     dupfinder.results = dupFindera(tree, salmonids = grep('huc', salmonid.tips, invert = T, value = T))
    test1 = is.monophyletic(drop.tip(tree, dupfinder.results$beta), dupfinder.results$alpha)
     test2 = is.monophyletic(drop.tip(tree, dupfinder.results$beta), grep('huc', dupfinder.results$alpha, invert
= T, value = T))
     return(identical(test1,test2)  & test1 == T)
      }
  dupfinder.results = dupFindera(tree) # dupfinder function adapted from Varadharajan et al, 2018
  if(length(dupfinder.results)>0){
    cat('Normal dup tree\n')
    test1 = is.monophyletic(drop.tip(tree, dupfinder.results$beta), dupfinder.results$alpha)
    test2 = is.monophyletic(drop.tip(tree, dupfinder.results$beta), grep('huc', dupfinder.results$alpha, invert
= T, value = T))
    return(identical(test1,test2)  & test1 == T)
  }
  if(length(dupfinder.results)==0){
    print('No dup')
    is.monophyletic(tree, grep('huc', tree$tip.label[substr(tree$tip.label, 1, 3) %in% salmonid.tips], invert =
T, value = T))
  }
}

#simple function to test for duplicate branching and SSR type.
lore_testing=function(tree){
  tree2<-read.tree(file = tree)
  if(length(grep('esox', tree2$tip.label, value = T))>0){
```

```
    test3<-root(tree2,outgroup = grep('esox', tree2$tip.label, value = T ), resolve.root = TRUE)
  }else if (length(grep('thym_chr_a', tree2$tip.label, value = T))>0){
    test3<-root(tree2,outgroup = grep('thym_chr_a', tree2$tip.label, value = T ), resolve.root = TRUE)
  }else if (length(grep('thym_chr_b', tree2$tip.label, value = T))>0){
    test3<-root(tree2,outgroup = grep('thym_chr_b', tree2$tip.label, value = T ), resolve.root = TRUE)
  #}else if (length(grep('huc', tree2$tip.label, value = T))==1){
    #test3<-root(tree2,outgroup = grep('huc', tree2$tip.label, value = T ), resolve.root = TRUE)
      }else {
    test3<-midpoint(tree2, node.labels = "label")}
  check.topologya(test3)
plot.coldupsa(tr = test3)
}

#standard looping - works good
files <- list.files(path="D:/trees/", pattern="*.treefile", full.names=TRUE, recursive=FALSE)
sink('ssr3_chra_chrb_10nodes.txt')
pdf("ssr3_chra_chrb_10nodes.txt ", width = 12, height = 12, paper = "a4r")
par(mfrow=c(2,1))
for(i in 1:length(files)){
  file<-cat(files[i],('\n'))
  lore_testing(files[i])
cat ('\n')}
sink()
dev.off()

#######get coordinates for salmon chromosomes from a MAF file (Bash script)#################
grep ssaXX.ssaXX mult11.maf | cut -c 1-55 | sort -n -k 3 >mult11_ssal_sorted_coordinates.txt

#############################################################
Retrieve genes from different SSR blocks and AORe regions in salmon genome (Rscript)
###############################################################
library(tidyverse)
library(openxlsx)
library(readxl)

##load data
aore_regions<-read_excel("aore_boundaries.xlsx")
aore_regions<-as.data.frame(apply(aore_regions,2,function(x)gsub('\\s+', '',x))) ### remove any spaces in
the whole data

lore_regions<-read_excel("lore_boundaries.xlsx")
lore_regions<-as.data.frame(apply(lore_regions,2,function(x)gsub('\\s+', '',x)))

chr_id_replace<-read_excel("ncbi_cigene_mapping.xlsx")
ssal_gene_pos<- read_tsv("Ssal_genePos.tsv")

duplicates<-read.table("duplicates_list.txt",header = T)
dup1<- duplicates$gene1
dup2<- duplicates$gene2
dups_list<- as.data.frame(c(dup1,dup2))
dups_list$`c(dup1, dup2)`<-as.numeric(dups_list$`c(dup1, dup2)`)

###change gene names to cigene
salmon_genes<-merge(ssal_gene_pos,chr_id_replace,by.x = c("seqname"), by.y = c("ncbi"),
      all.x = T, all.y = F) %>% na.omit(T) %>% select(cigene,start,end,geneID) %>% arrange(cigene,start)
```

```
salmon_genes$start <- as.numeric(salmon_genes$start)
salmon_genes$end <- as.numeric(salmon_genes$end)

###### function to get genes within aore boundaries
try1<-lapply(1:nrow(aore_regions),function(i){
  ref_chro <- as.character(aore_regions$chr[i])
  ref_start <- as.numeric(aore_regions$start[i])
  ref_end <- as.numeric(aore_regions$end[i])
  filter(salmon_genes,cigene == ref_chro,start > ref_start & start < ref_end)
  })

aore_genes<-bind_rows(try1)
aore_genes['redip'] = "AORe"

#### function against lore regions
lore_redip <- filter(lore_regions, redip == "SSR1") ## set SSR1, SSR2. SSR3,SSR4,SSR5

try2<-lapply(1:nrow(lore_redip),function(i){
  ref_chro <- as.character(lore_redip$chr[i])
  ref_start <- as.numeric(lore_redip$start[i])
  ref_end <- as.numeric(lore_redip$end[i])
  filter(salmon_genes,cigene == ref_chro,start > ref_start & start < ref_end)
})

SSR1_genes<-bind_rows(try2)
SSR1_genes['redip'] = "SSR1"
SSR1_genes$geneID = as.numeric(SSR1_genes$geneID)

SSR2_genes<-bind_rows(try2)
SSR2_genes['redip'] = "SSR2"

SSR3_genes<-bind_rows(try2)
SSR3_genes['redip'] = "SSR3"

SSR4_genes<-bind_rows(try2)
SSR4_genes['redip'] = "SSR4"

SSR5_genes<-bind_rows(try2)
SSR5_genes['redip'] = "SSR5"

### compare it against the duplicates database from Bertolotti et al
aore_dups<-as.data.frame(intersect(dups_list$`c(dup1, dup2)`, aore_genes$geneID))
ssr1_dups<-as.data.frame(intersect(dups_list$`c(dup1, dup2)`, SSR1_genes$geneID))
ssr2_dups<-as.data.frame(intersect(dups_list$`c(dup1, dup2)`, SSR2_genes$geneID))
ssr3_dups<-intersect(dups_list$`c(dup1, dup2)`, SSR3_genes$geneID)
ssr4_dups<-intersect(dups_list$`c(dup1, dup2)`, SSR4_genes$geneID)
ssr5_dups<-intersect(dups_list$`c(dup1, dup2)`, SSR5_genes$geneID)

ssr_345_dups<- as.data.frame(c(ssr3_dups,ssr4_dups,ssr5_dups)) ### merge all ssr345 into one dataframe

gene_annotation<-read_csv("genes_products.csv")

aore_dups_annotation<-left_join(aore_dups,gene_annotation)
ssr1_dups_annotation<-left_join(ssr1_dups,gene_annotation)
```

```
ssr2_dups_annotation<-left_join(ssr2_dups,gene_annotation)
ssr345_dups_annotation<-left_join(ssr345_dups,gene_annotation)
```

```
##########################################################################################
```

Get unique and shared GOslim terms from different rediploidization categories (R script)

````
###########################################################################
###load data

daore<-read_excel("GO_slim_to_plot.xlsx", sheet = "AORE_ohno")
daore2<-daore %>% select(Term, Count)
daore2['redip']='AORe-ohnologue'

dssr1<-read_excel("GO_slim_to_plot.xlsx", sheet = "SSR1_ohno")
dssr12<-dssr1 %>% select(Term, Count)
dssr12['redip']='SSR1-ohnologue'

dssr2<-read_excel("GO_slim_to_plot.xlsx", sheet = "SSR2_ohno")
dssr22<-dssr2 %>% select(Term, Count)
dssr22['redip']='SSR2-ohnologue'

dssr345<-read_excel("GO_slim_to_plot.xlsx", sheet = "SSR345_ohno")
dssr3452<-dssr345 %>% select(Term, Count)
dssr3452['redip']='SSR345-ohnologue'

saore<-read_excel("GO_slim_to_plot.xlsx", sheet = "AORE_sing")
saore2<-saore %>% select(Term, Count)
saore2['redip']='AORe-singleton'
unique_singletons<- as.vector(saore2$Term)

####pool data and aggregate

pool1<-rbind(saore2,daore2,dssr12,dssr22,dssr3452)

upset_data<-aggregate(redip ~ Term, unique(pool1), paste, collapse = "&")  ## aggregate by GO terms

datapool<-aggregate(redip ~ Term, unique(pool1), paste, collapse = ",")   ## aggregate by GO terms

GO_slim_unique_shared<- datapool %>% arrange(redip)


##########################################################################################
````

R script to parse orthogroups and get dN/dS values for selected nodes

````
#####################################################################################
---
title: "Identify branches of interest in aBSREL results"
output:
  html_document:
    toc: yes
    toc_float: yes
    code_folding: hide
editor_options:
  chunk_output_type: console
---
```{r setup, include=FALSE}
````
````

````
knitr::opts_chunk$set(echo = TRUE)
library(tidyverse)
library(ape)
library(jsonlite)
```

```{r load_data}
source("getEdgesToClade.R")
# extract trees from json files:
extractTreesFromJSON <- function(jsonpath){
  tibble( file = dir(jsonpath,full.names = T,pattern = ".json$")) %>%
    mutate( data = map( file, read_json)) %>%
    mutate( jsonTree = map( data, ~ read.tree(text = paste0(.x$input$trees$`0`,";")) )) %>%
    mutate( OG = sub(".fasta.ABSREL.json","",basename(file))) %>%
    with( setNames(jsonTree,OG))
}

resultPath <- "~/Dropbox/REWIRED project/received files/from_manu/aBSREL results for Manu article"

ssr1Trees <- extractTreesFromJSON(file.path(resultPath,"ssr1_json"))
ssr2Trees <- extractTreesFromJSON(file.path(resultPath,"ssr2_json"))
aoreTrees <- extractTreesFromJSON(file.path(resultPath,"aore_json"))

spcTree <- read.tree("data/from_ortho_pipeline/SpeciesTree_rooted_node_labels.txt")
# define some useful constants
N6spcs <- c("Stru","Ssal","Omyk","Okis","Salp")
N5spcs <- c(N6spcs,"Hhuc")
N4spcs <- c(N5spcs,"Tthy")

```
#### Species tree
```{r}
plot.phylo(spcTree, show.node.label = T,use.edge.length = F)
```

#### Identifying clades
The aim is to identify the branches of the gene-trees that lead to specific clades defined by a set of species.
The algorithm I use here identifies edges in the tree where the all children of the child node are in the set of clade species, with no copies, but the same is not true for the parent node.
#### branches of interest
##### AORE - Ortholog resolution at salmonid common ancestor
Branches of interest are:
* (blue) the branches going from duplication node to the two N4 clades (All salmonids)
* (red) the branches going to the two N6 clades (All salmonids except Tthy and Hhuc)

###### Example

```{r}
OG="OG1v0000101.121"
tree = aoreTrees[[OG]]
edgeColors <- rep("black",Nedge(tree))
edgeColors[getEdgesToClade(tree,N6spcs)] <- "red"
edgeColors[getEdgesToClade(tree,N4spcs)] <- "blue"
plot(tree, use.edge.length = F,edge.color = edgeColors,edge.width = 2, main=OG)
```
````
````

###### stats
How many clades do we identify in each tree? Do they overlap? Are the clades rooted in internal nodes (as opposed to tips)?
```{r}
trees = aoreTrees

N6cladeEdges <- lapply(trees, getEdgesToClade, cladeSpcs = N6spcs)
N4cladeEdges <- lapply(trees, getEdgesToClade, cladeSpcs = N4spcs)
N_N6clades <- sapply(N6cladeEdges,length)
N_N4clades <- sapply(N4cladeEdges,length)
overlap <- sapply(mapply(FUN = intersect,x=N6cladeEdges,y=N4cladeEdges),length)
table(N_N6clades,N_N4clades,overlap=overlap>0)
edges2nodeNames <- function(edges,tree){
  nodes <- tree$edge[edges,2]
  c(tree$tip.label,tree$node.label)[nodes]
}

N6nodeNames <- mapply( edges = N6cladeEdges, tree=trees, FUN=edges2nodeNames)
N4nodeNames <- mapply( edges = N4cladeEdges, tree=trees, FUN=edges2nodeNames)
allN6nodesInternal <- sapply(N6nodeNames,function(x) all(grepl("^Node",x)))
allN4nodesInternal <- sapply(N4nodeNames,function(x) all(grepl("^Node",x)))

goodTrees <- N_N6clades==2 & N_N4clades==2 & overlap==0

table(allN6nodesInternal,goodTrees)

aore_goodTrees <- N_N6clades==2 & N_N4clades==2 & overlap==0 & allN6nodesInternal &
allN4nodesInternal

```
Of the `r length(trees)` AORE trees, `r sum(aore_goodTrees)` trees have exactly two N4 clades and two N6 clades that do not overlap.
###### Bad tree #1
Only one inner N4 clade:

```{r}
# which(N_N6clades==1 & N_N4clades==2 & overlap==0)
OG="OG1v0011980.15"
tree = trees[[OG]]

edgeColors <- rep("black",Nedge(tree))
edgeColors[getEdgesToClade(tree,N6spcs)] <- "red"
edgeColors[getEdgesToClade(tree,N4spcs)] <- "blue"

plot(tree, use.edge.length = F,edge.color = edgeColors,edge.width = 2,main=OG)
```

###### Bad tree #2
One clade is missing both Tthy and Hhuc, therefore N4 clade is same as N6 clade:
```{r}
# which(N_N6clades==2 & N_N4clades==2 & overlap>0)
OG="OG1v0014017.13"
tree = trees[[OG]]

edgeColors <- rep("black",Nedge(tree))
```

edgeColors[getEdgesToClade(tree,N6spcs)] <- "red"
edgeColors[getEdgesToClade(tree,N4spcs)] <- "blue"
edgeColors[intersect(getEdgesToClade(tree,N4spcs),getEdgesToClade(tree,N6spcs))] <- "purple"
plot(tree, use.edge.length = F,edge.color = edgeColors,edge.width = 2,main=OG)
```

### SSR1 - Ortholog resolution after Tthy split
Branches of interest are:
* (blue) the branches going from duplication node to the two N5 clades (All salmonids except Tthy)
* (red) the branches going to the two N6 clades (All salmonids except Tthy and Hhuc)
#### Example
```{r}
tree = ssr1Trees$OG1v0000520.54

edgeColors <- rep("black",Nedge(tree))
edgeColors[getEdgesToClade(tree,N6spcs)] <- "red"
edgeColors[getEdgesToClade(tree,N5spcs)] <- "blue"

plot(tree, use.edge.length = F,edge.color = edgeColors,edge.width = 2, main="OG1v0000520.54")
```

#### stats
How many clades do we identify in each tree?
```{r}
trees = ssr1Trees

N6cladeEdges <- lapply(trees, getEdgesToClade, cladeSpcs = N6spcs)
N5cladeEdges <- lapply(trees, getEdgesToClade, cladeSpcs = N5spcs)
N_N6clades <- sapply(N6cladeEdges,length)
N_N5clades <- sapply(N5cladeEdges,length)

overlap <- sapply(mapply(FUN = intersect,x=N6cladeEdges,y=N5cladeEdges),length)

table(N_N6clades,N_N5clades,overlap=overlap>0)


N6nodeNames <- mapply( edges = N6cladeEdges, tree=trees, FUN=edges2nodeNames)
N5nodeNames <- mapply( edges = N5cladeEdges, tree=trees, FUN=edges2nodeNames)

allN6nodesInternal <- sapply(N6nodeNames,function(x) all(grepl("^Node",x)))
allN5nodesInternal <- sapply(N5nodeNames,function(x) all(grepl("^Node",x)))

goodTrees <- N_N6clades==2 & N_N5clades==2 & overlap==0

ssr1_goodTrees <- N_N6clades==2 & N_N5clades==2 & overlap==0 & allN6nodesInternal &
allN5nodesInternal

```
Of the `r length(ssr1Trees)` ssr1 trees, `r sum(ssr1_goodTrees)` trees have exactly two N5 clades and two N6 clades that do not overlap.
#### Bad tree #1

```{r}
### which(N_ssr1N6clades==9 & N_ssr1N5clades==12)

tree = ssr1Trees$OG1v0000994.33

edgeColors <- rep("black",Nedge(tree))
edgeColors[getEdgesToClade(tree,N6spcs)] <- "red"
edgeColors[getEdgesToClade(tree,N5spcs)] <- "blue"

plot(tree, use.edge.length = F,edge.color = edgeColors,edge.width = 2,main="OG1v0000994.33")
```

Completely messed up...
#### Bad tree #2
Contains duplicate Hhuc gene:
```{r}
### which(N_ssr1N6clades==2 & N_ssr1N5clades==3 & ssr1_overlap==0)
OG = "OG1v0004789.19"
tree = ssr1Trees[[OG]]

edgeColors <- rep("black",Nedge(tree))
edgeColors[getEdgesToClade(tree,N6spcs)] <- "red"
edgeColors[getEdgesToClade(tree,N5spcs)] <- "blue"

plot(tree, use.edge.length = F,edge.color = edgeColors,edge.width = 2,main=OG)
```
### SSR2 - Ortholog resolution after Hhuc split
Branches of interest are:
* (red) the branches going from duplication node to the two N6 clades (All salmonids except Tthy and Hhuc)

#### Example
```{r}
### filter(OGtbl, geneID=="105019051")$N0
OG="OG1v0000596.55"
tree = ssr2Trees[[OG]]
edgeColors <- rep("black",Nedge(tree))
edgeColors[getEdgesToClade(tree,N6spcs)] <- "red"
plot(tree, use.edge.length = F,edge.color = edgeColors,edge.width = 2, main=OG)
```

#### stats
How many clades do we identify in each tree? Do they overlap? Are the clades rooted in internal nodes (as opposed to tips)?
```{r}
trees = ssr2Trees

N6cladeEdges <- lapply(trees, getEdgesToClade, cladeSpcs = N6spcs)
N_N6clades <- sapply(N6cladeEdges,length)
table(N_N6clades)
N6nodeNames <- mapply( edges = N6cladeEdges, tree=trees, FUN=edges2nodeNames)
allN6nodesInternal <- sapply(N6nodeNames,function(x) all(grepl("^Node",x)))
goodTrees <- N_N6clades==2
table(allN6nodesInternal,goodTrees)
ssr2_goodTrees <- N_N6clades==2 & allN6nodesInternal

```
```

Of the `r length(trees)` SSR2 trees, `r sum(ssr2_goodTrees)` trees have exactly two N6 clades which are not tip nodes.
###### Bad tree #1
Salp duplicates form a clade:

```{r}
which(N_N6clades==4)
OG="OG1v0000981.48"
tree = trees[[OG]]

edgeColors <- rep("black",Nedge(tree))
edgeColors[getEdgesToClade(tree,N6spcs)] <- "red"

plot(tree, use.edge.length = F,edge.color = edgeColors,edge.width = 2,main=OG)
```

###### Bad tree #2

```{r}
# which(N_N6clades==4)
OG="OG1v0006983.18"
tree = trees[[OG]]
edgeColors <- rep("black",Nedge(tree))
edgeColors[getEdgesToClade(tree,N6spcs)] <- "red"
plot(tree, use.edge.length = F,edge.color = edgeColors,edge.width = 2,main=OG)
```

#### CSV files
Combine all the csv files for the "good" trees and add a column identifying the nodes of interest.
```{r}

getCladeNodeNames <- function(tree,cladeSpcs){
  edges <- getEdgesToClade(tree,cladeSpcs)
  edges2nodeNames(edges,tree)
}

nodeTbl <- bind_rows(
  .id="oreType",
  aore = tibble(OG = names(aore_goodTrees)[aore_goodTrees]) %>%
    mutate( N6 = map(OG, function(OG){getCladeNodeNames(tree=aoreTrees[[OG]], cladeSpcs =
N6spcs)})) %>%
    mutate( N4 = map(OG, function(OG){getCladeNodeNames(tree=aoreTrees[[OG]], cladeSpcs =
N4spcs)})) %>%
    gather(key = "clade",value = "node",N6,N4) %>%
    unnest(node),
  ssr1 = tibble(OG = names(ssr1_goodTrees)[ssr1_goodTrees]) %>%
    mutate( N6 = map(OG, function(OG){getCladeNodeNames(tree=ssr1Trees[[OG]], cladeSpcs =
N6spcs)})) %>%
    mutate( N5 = map(OG, function(OG){getCladeNodeNames(tree=ssr1Trees[[OG]], cladeSpcs =
N5spcs)})) %>%
    gather(key = "clade",value = "node",N6,N5) %>%
    unnest(node),
  ssr2 = tibble(OG = names(ssr2_goodTrees)[ssr2_goodTrees]) %>%
    mutate( N6 = map(OG, function(OG){getCladeNodeNames(tree=ssr2Trees[[OG]], cladeSpcs =
N6spcs)})) %>%
```

````
    gather(key = "clade",value = "node",N6) %>%
    unnest(node)
)
combinedCSVtable <-
  nodeTbl %>%
  select(oreType, OG) %>%
  distinct() %>%
  mutate( filename=file.path(resultPath,paste0(oreType,"_json"),
                  paste0(OG,".fasta.ABSREL.json.csv")) ) %>%
  mutate( tbl = map(filename,read_csv,col_types = cols())) %>%
  select(-filename) %>%
  unnest(tbl) %>%
  left_join(nodeTbl, by=c("oreType","OG","node"))

write_csv(combinedCSVtable,"combinedCSVtable.csv")
```
#### plot the dN/dS values for for the identified clades
```{r}
combinedCSVtable %>%
  filter( !is.na(clade)) %>%
  filter( baseline_omega < 200) %>%
  ggplot( aes(x=clade,y=baseline_omega)) + geom_boxplot() + facet_grid( . ~ oreType) +
  coord_cartesian(ylim=c(0,2))
```
````

**Supplementary Methods 3: SSR trees mapped to the brown trout genome**

The chromosome level genome of brown trout (NCBI accession: GCA_901001165.1) was used to explore the impact of genome assembly on the mapping of SSR tree topologies (Supplementary Fig. 11-12). The goal was to answer whether the physical interspersing of distinct SSR phylogenetic topologies along Atlantic salmon chromosomes was caused by assembly artefacts. For this test, we selected the ohnologue block represented by Atlantic salmon chromosome arms 03 and 06 (Ssa03-06) owing to the large number of recovered phylogenetic trees. Brown trout sequences homologous to Ssa03 and Ssa06 were identified by BLASTn as chromosomes 01 and 32 (St01, St32). Multispecies genome alignments accommodating these brown trout sequences were generated and processed using the methods described in the main text, except that the resultant tree topologies were visualized along St01 and St32 instead of Ssa03 and Ssa06 (Supplementary Fig. 11). In addition, a circos plot was generated to compare Ssa06 with its brown trout putative orthologue St01 to consider the impact of rearrangements across species (Supplementary Fig. 12).
````

#### Supplementary Figures

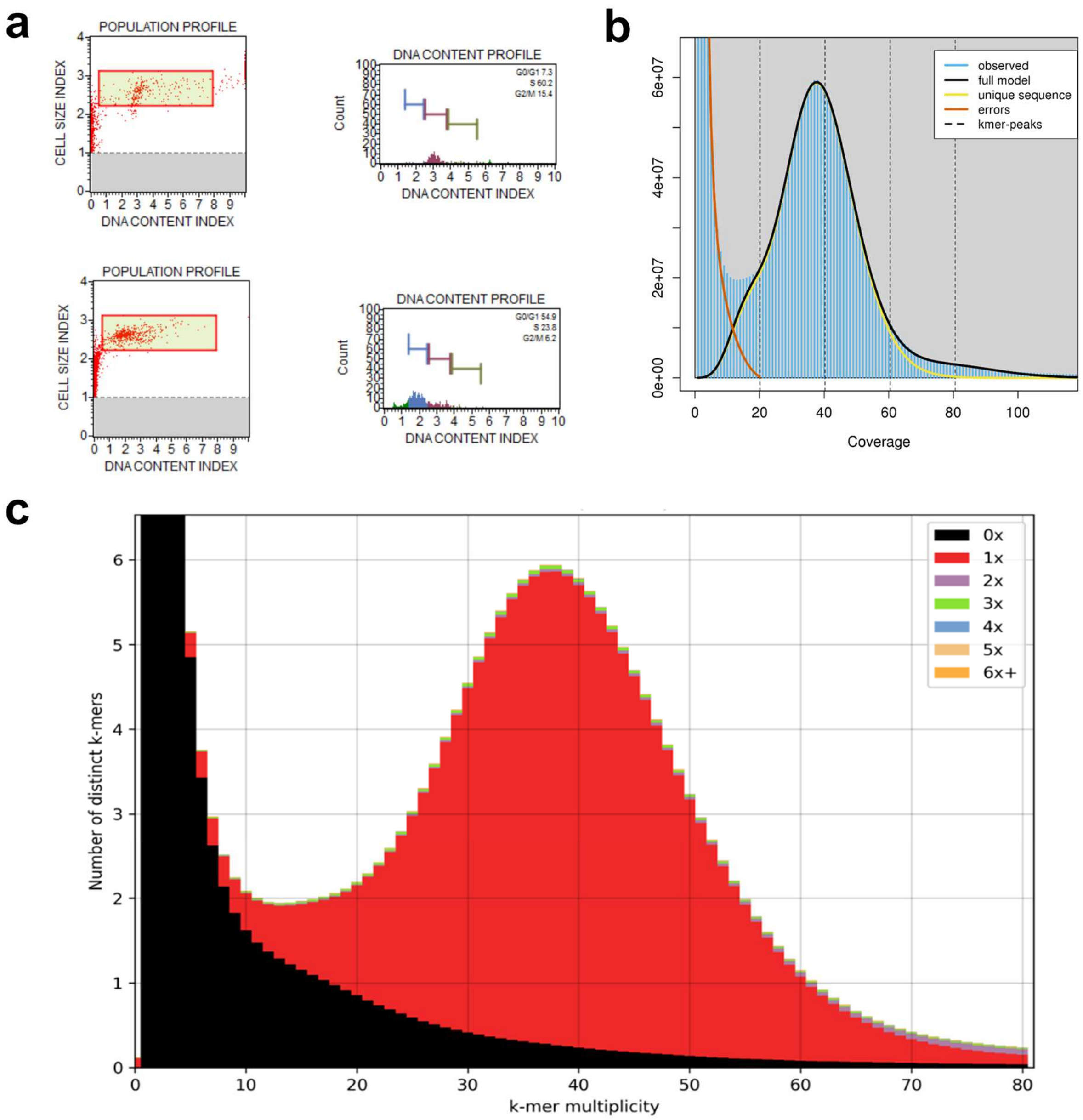


**Supplementary Fig. 1.** Huchen genome assembly validation. **a** Flow cytometry validation of haploidy in huchen. DNA content index for a diploid huchen individual (top panel) and for the sequenced haploid embryo (bottom panel). **b** Genomescope k-mer distribution for the huchen using paired-end sequence data (k-mer value: 101) **c** KAT plot for the gap-filled huchen assembly. The black region represents paired-end sequences that are not present in the assembly. The red region represents paired-end sequences used once in the assembly. The distribution of the red area indicates a homozygous assembly.

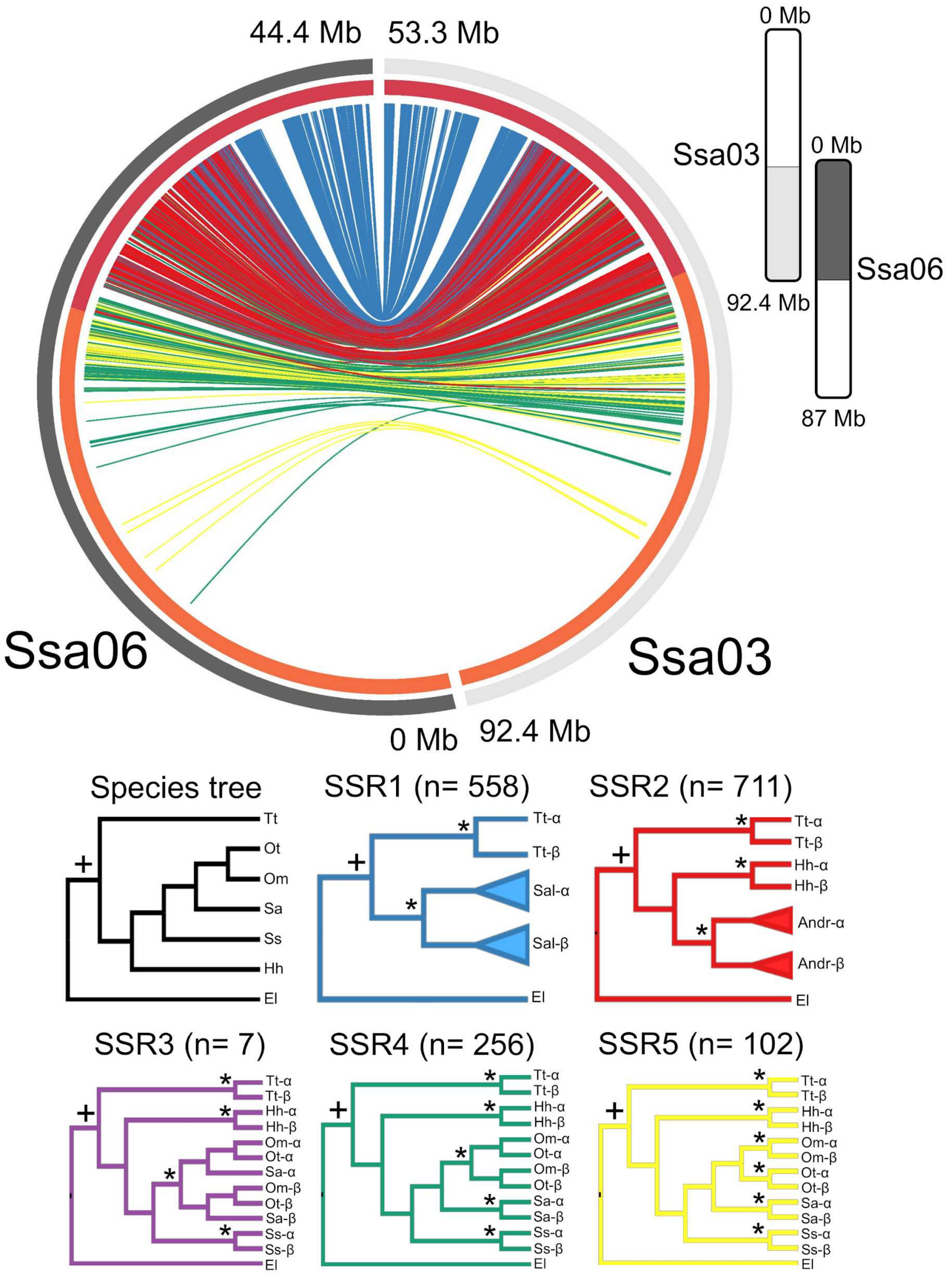


**Supplementary Fig. 2.** Mapping of reconstructed lineage-specific rediploidization histories for ohnologue regions Ssa03 and Ssa06 in the Atlantic salmon genome. A circos plot is shown at the top of the figure, where each sweeping band represents the location of ohnologue regions sampled in sequence alignments on Ssa03 and Ssa06. The panel immediately proximal to these bands shows the percent nucleotide sequence identity between the duplicated regions; orange = 90-95%, red = > 95%. The colour of each band represents the phylogenetic topology reconstructed for each sampled alignment with respect to trees presented below the circos plot based on matching colours (e.g. sweeping blue bands represent SSR1 topologies). We also provide the number (n) of sampled trees per defined rediploidization history.

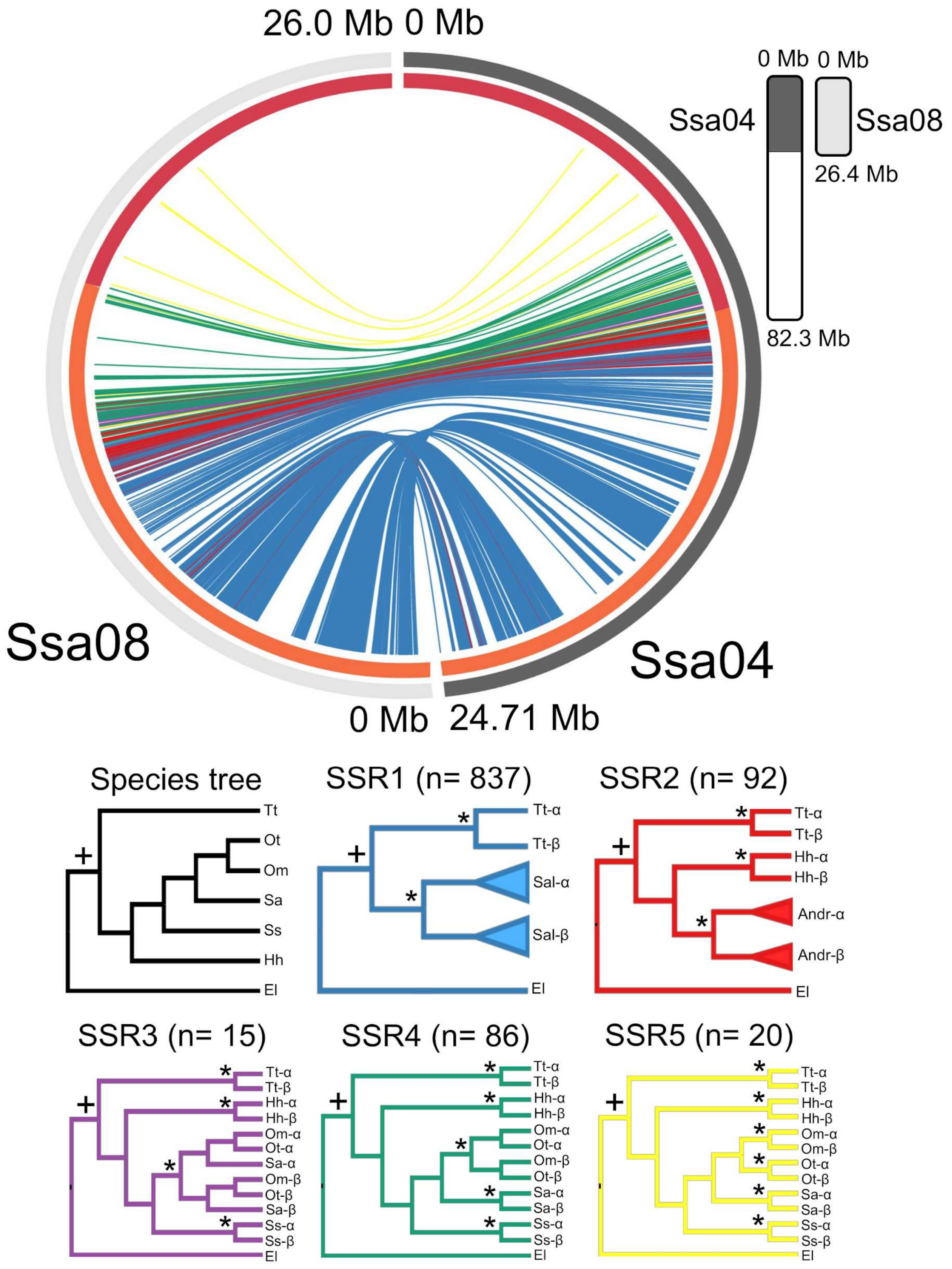


**Supplementary Fig. 3.** Mapping of reconstructed lineage-specific rediploidization histories for ohnologue regions Ssa04 and Ssa08 in the Atlantic salmon genome. All other details are as described in the Supplementary Fig. 2 legend.

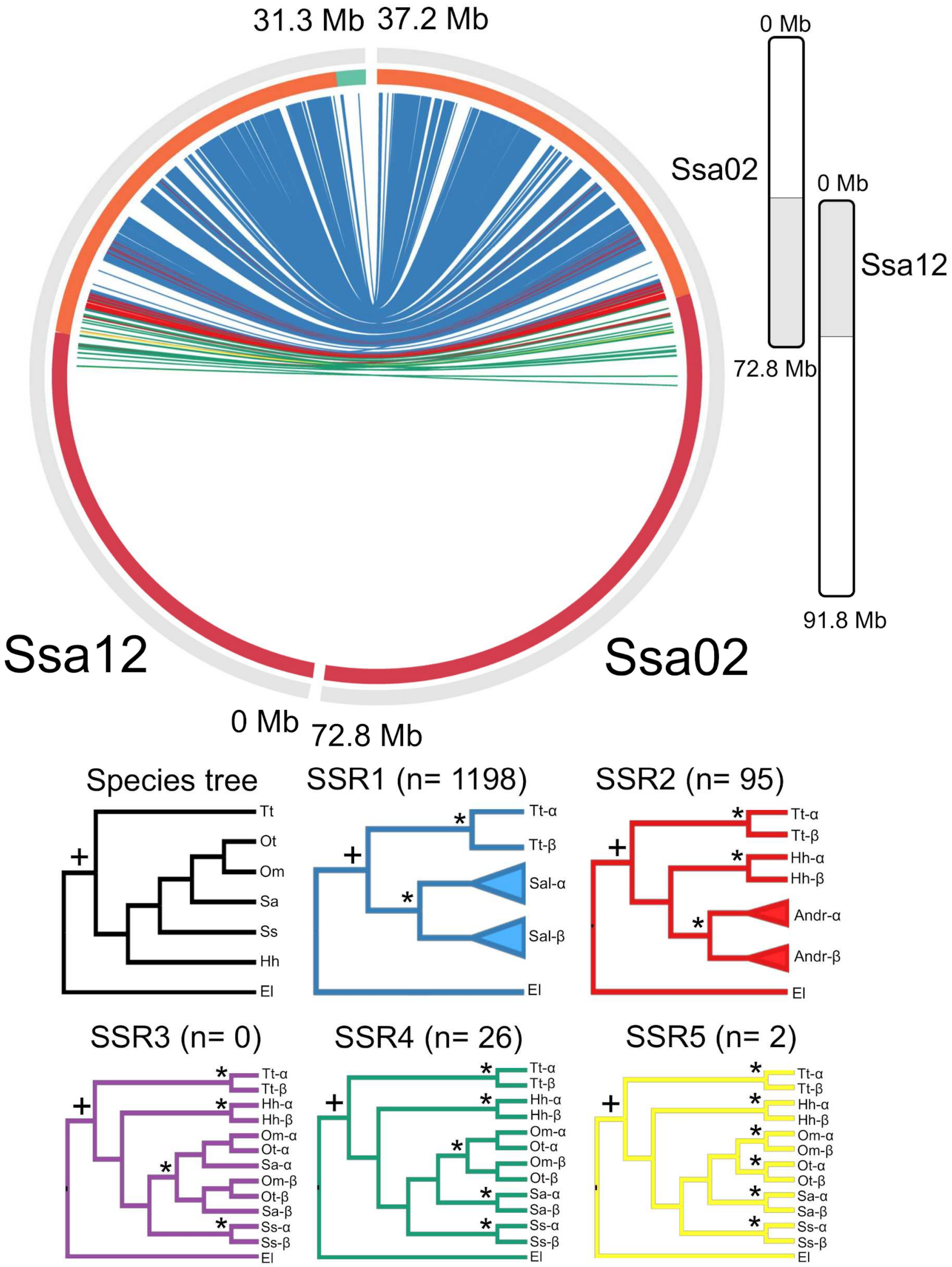


**Supplementary Fig. 4.** Mapping of reconstructed lineage-specific rediploidization histories for ohnologue regions Ssa02 and Ssa12 in the Atlantic salmon genome. All other details are as described in the Supplementary Fig. 2 legend.

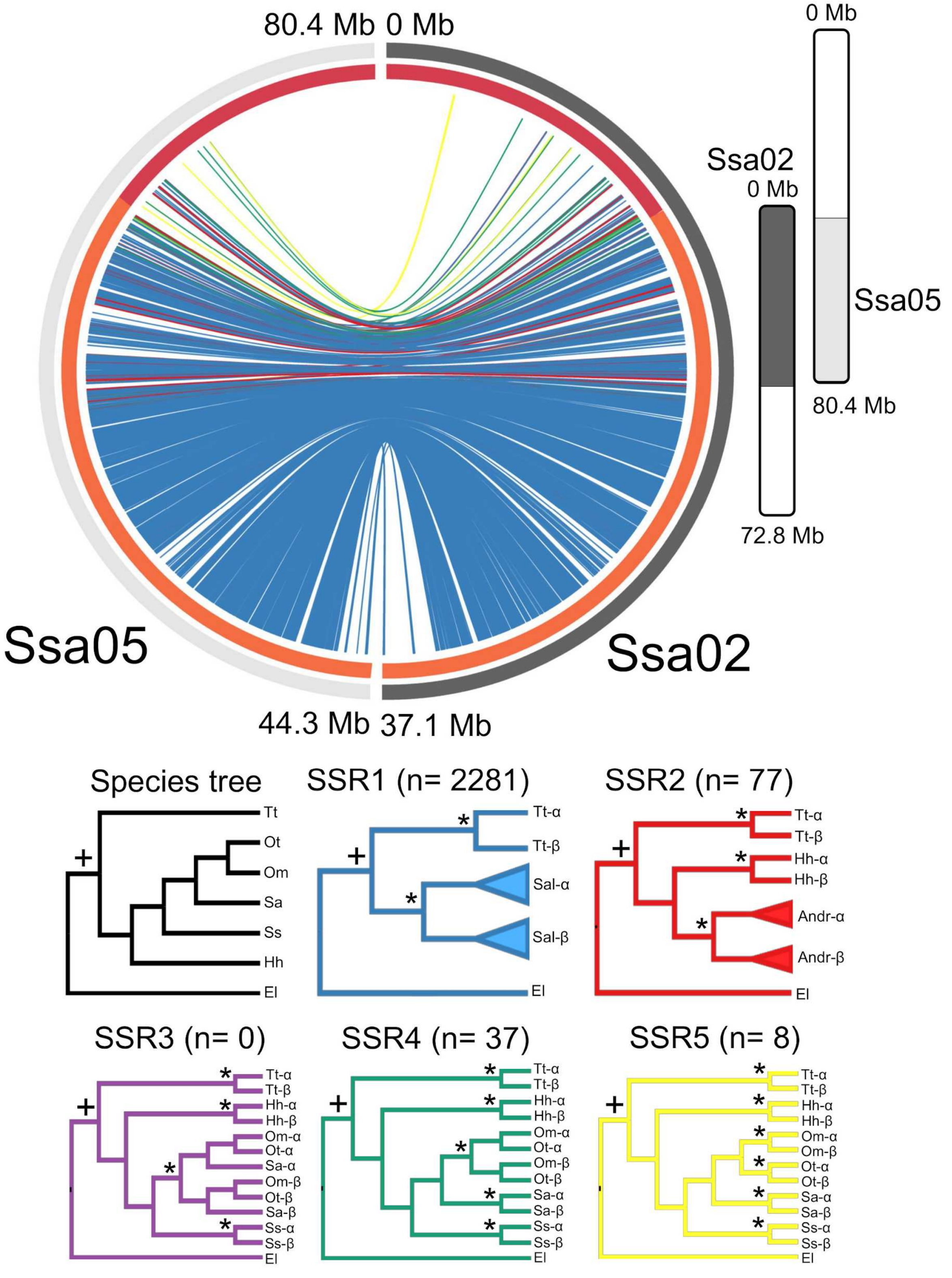


**Supplementary Fig. 5.** Mapping of reconstructed lineage-specific rediploidization histories for ohnologue regions Ssa02 and Ssa05 in the Atlantic salmon genome. All other details are as described in the Supplementary Fig. 2 legend.

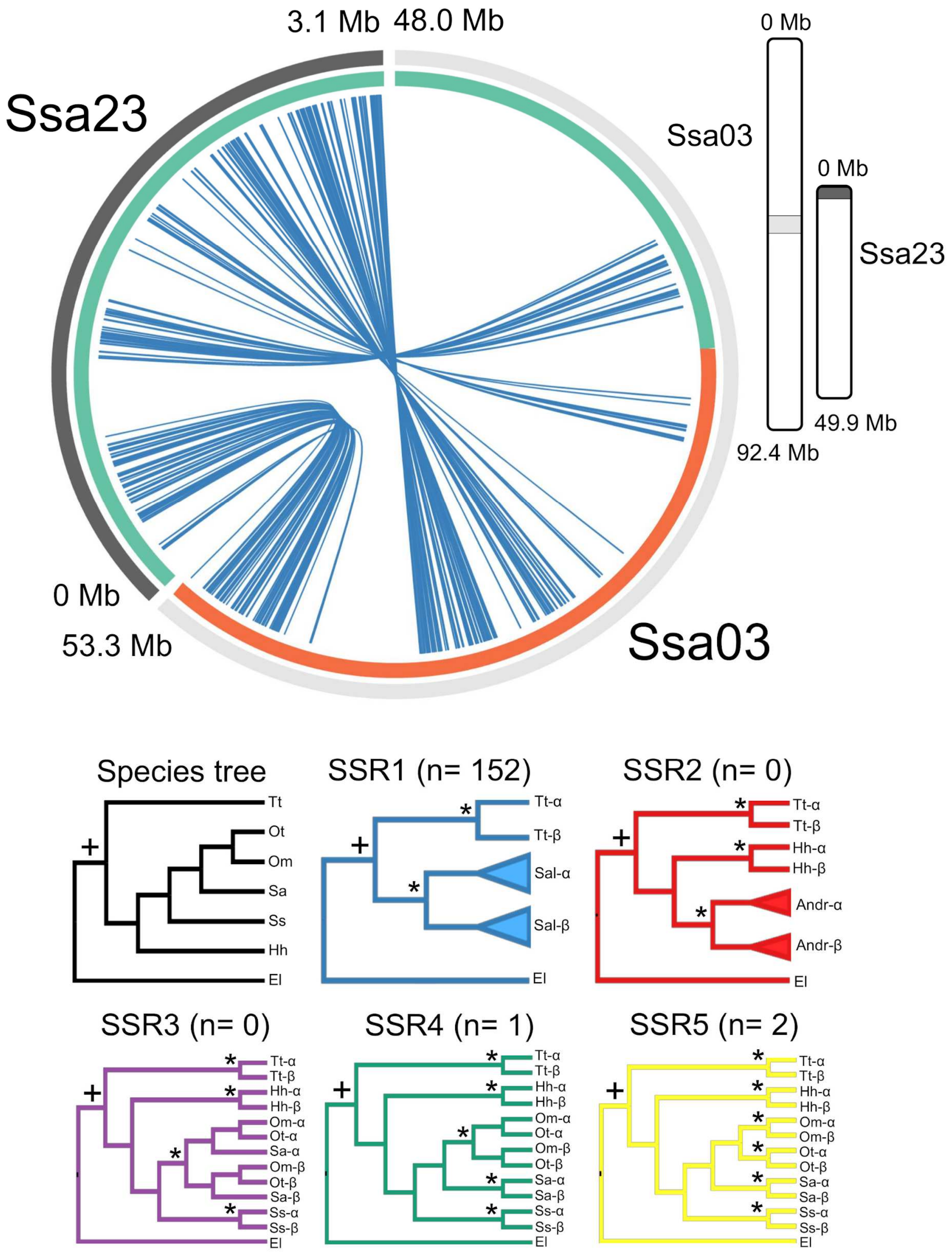


**Supplementary Fig. 6.** Mapping of reconstructed lineage-specific rediploidization histories for ohnologue regions Ssa03 and Ssa23 in the Atlantic salmon genome. All other details are as described in the Supplementary Fig. 2 legend.

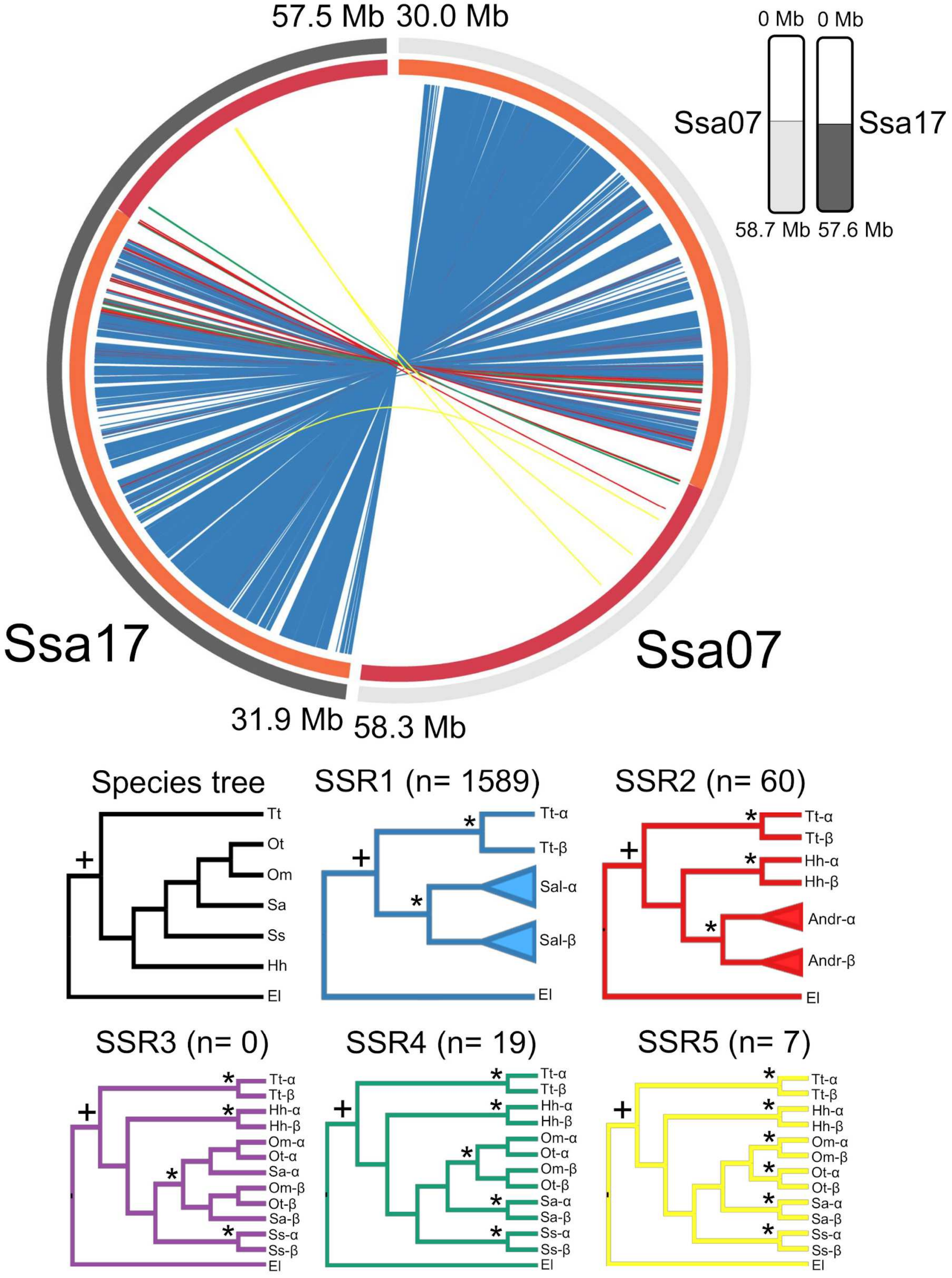


**Supplementary Fig. 7.** Mapping of reconstructed lineage-specific rediploidization histories for ohnologue regions Ssa07 and Ssa17 in the Atlantic salmon genome. All other details are as described in the Supplementary Fig. 2 legend.

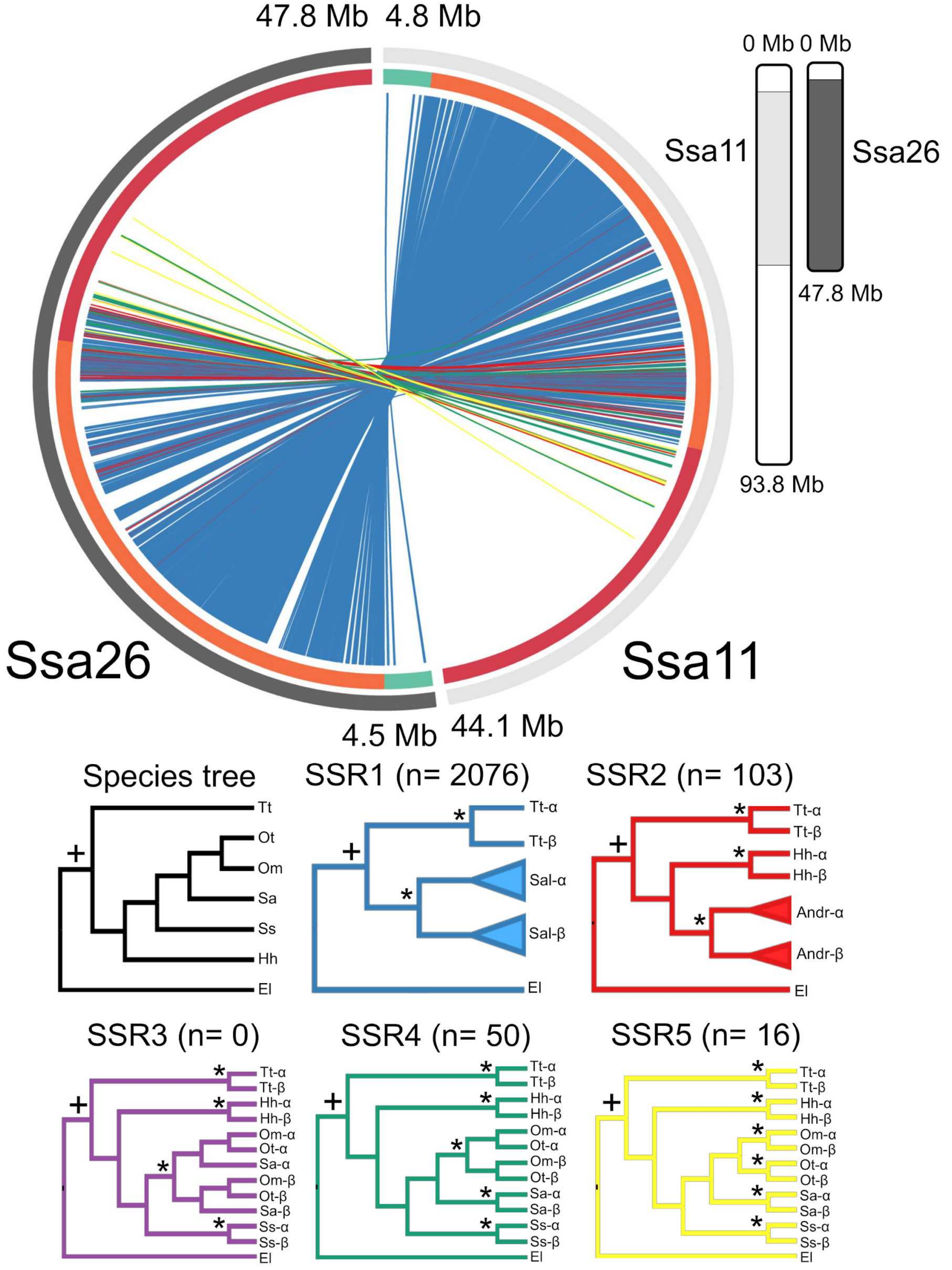


**Supplementary Fig. 8.** Mapping of reconstructed lineage-specific rediploidization histories for ohnologue regions Ssa11 and Ssa26 in the Atlantic salmon genome. All other details are as described in the Supplementary Fig. 2 legend.

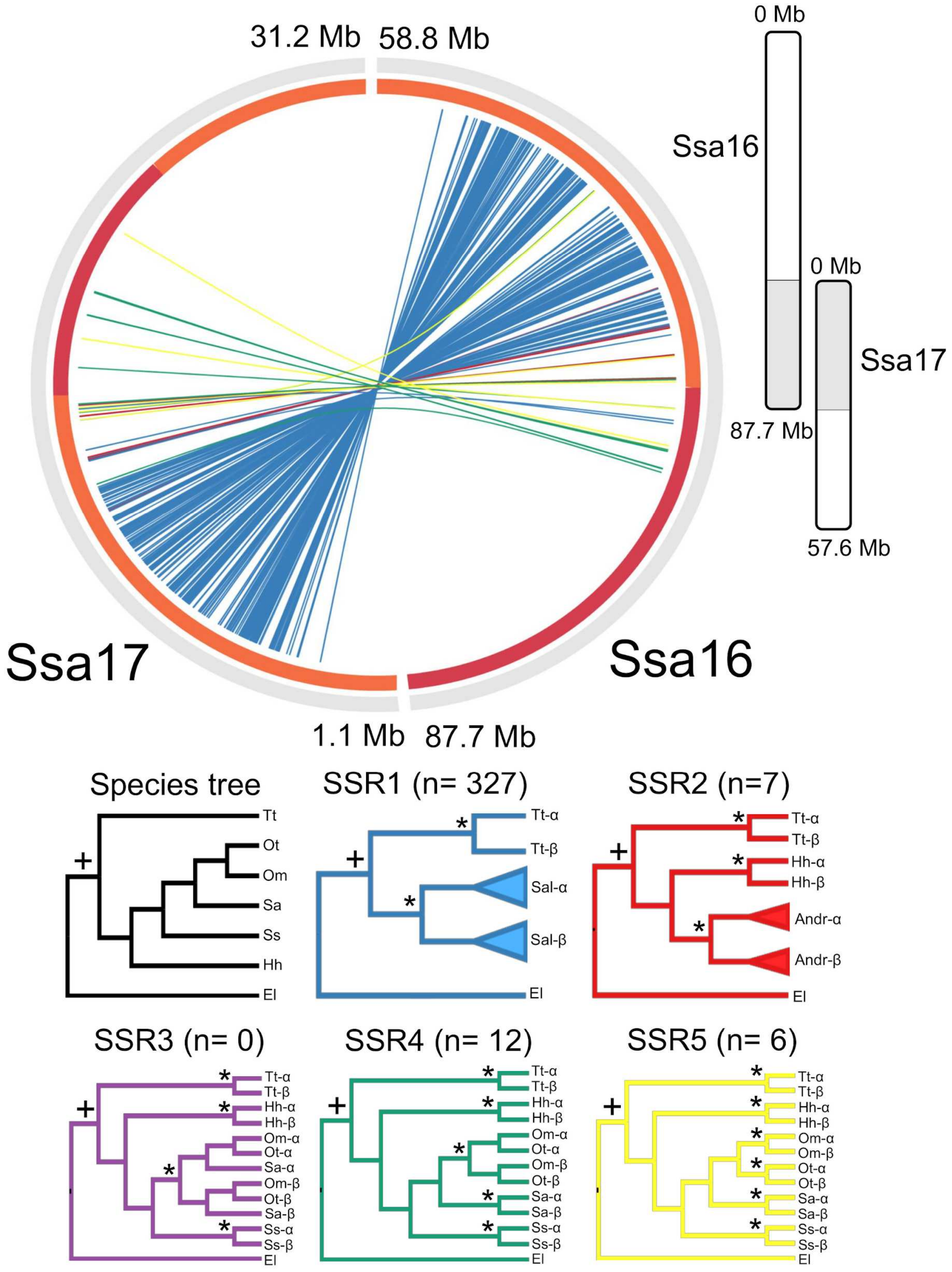


**Supplementary Fig. 9.** Mapping of reconstructed lineage-specific rediploidization histories for ohnologue regions Ssa16 and Ssa17 in the Atlantic salmon genome. All other details are as described in the Supplementary Fig. 2 legend.

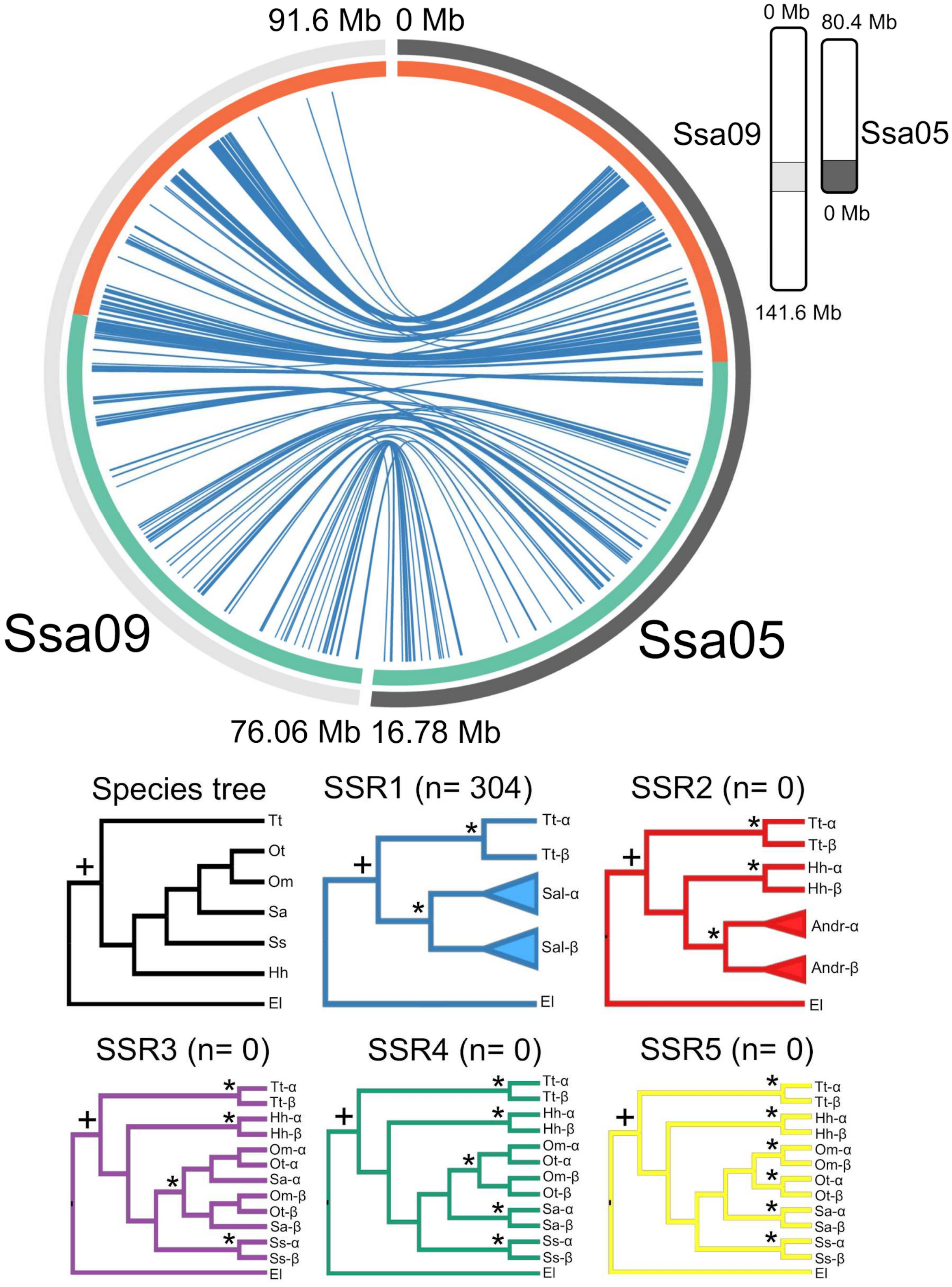


**Supplementary Fig. 10.** Mapping of reconstructed lineage-specific rediploidization histories for ohnologue regions Ssa05 and Ssa09 in the Atlantic salmon genome. All other details are as described in the Supplementary Fig. 2 legend.

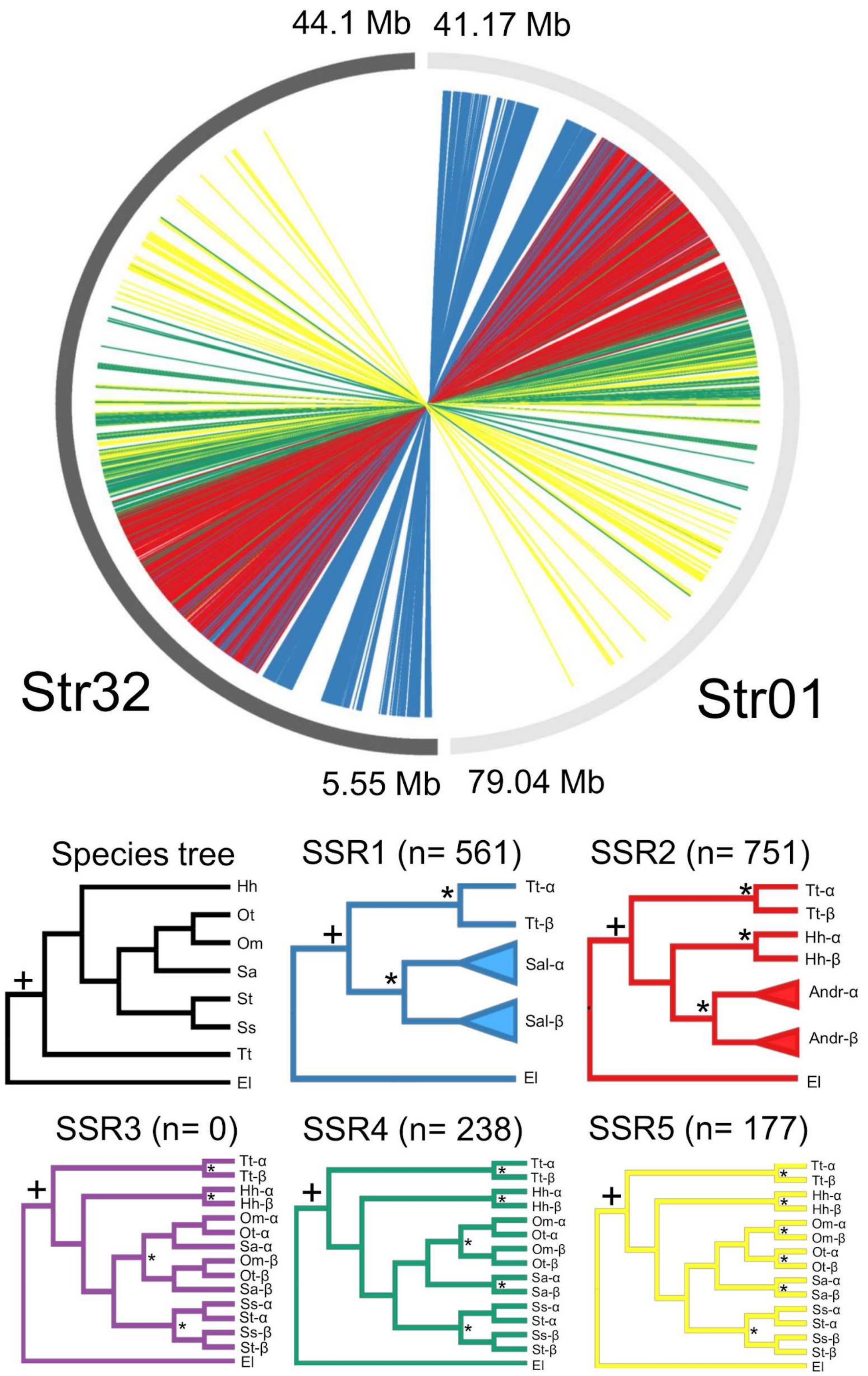


**Supplementary Fig. 11.** Mapping of reconstructed lineage-specific rediploidization histories for ohnologue regions Str01 and Str32 in the brown trout genome (orthologous to Ssa06 and Ssa03 in the Atlantic salmon genome, respectively). This figure demonstrates a similar intermixing of phylogenetic signals (i.e. different rediploidization histories) in the brown trout genome as observed for the Atlantic salmon genome (compare with Supplementary Fig. 3). All other details are as described in the Supplementary Fig. 2 legend.

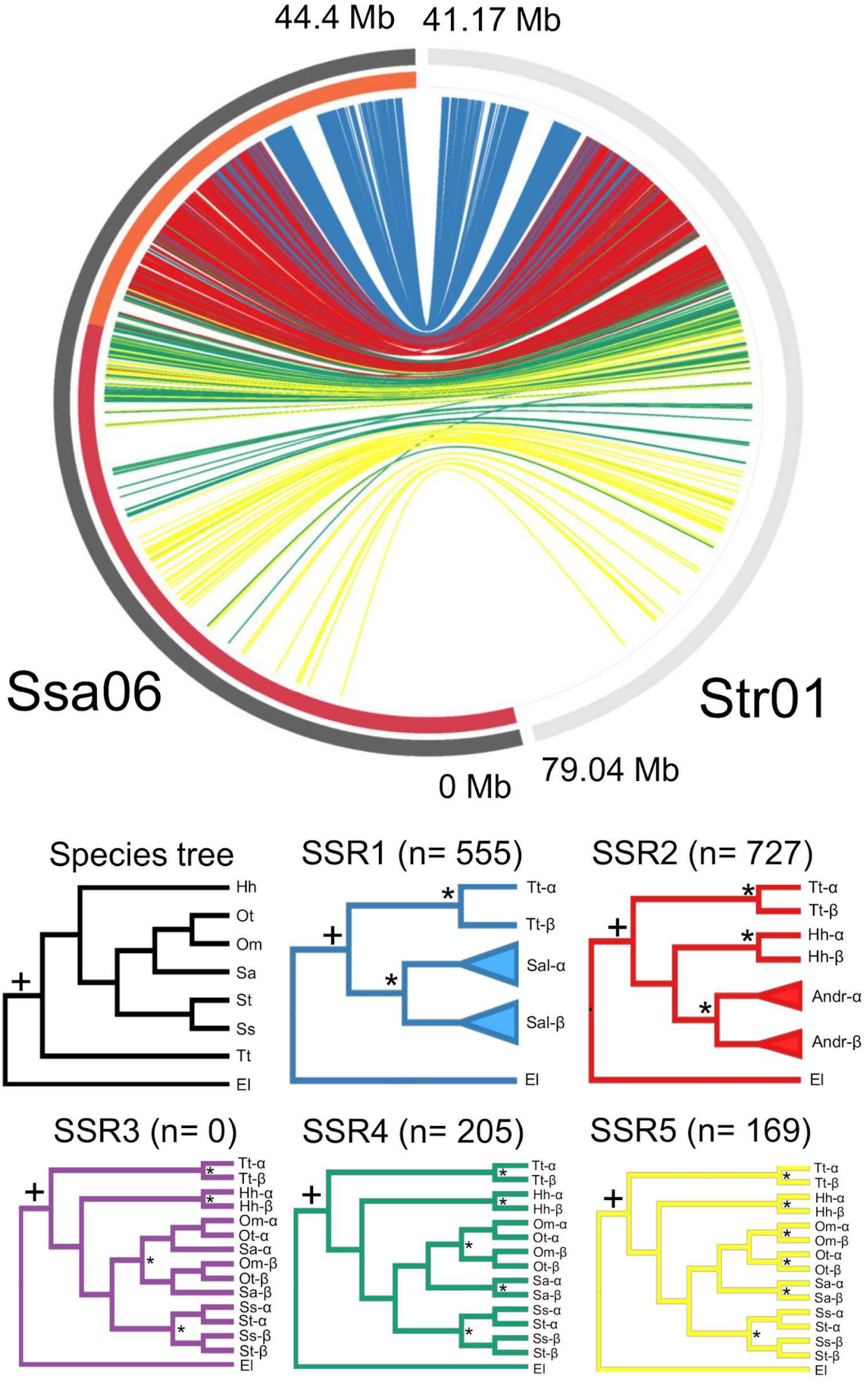


**Supplementary Fig. 12.** Mapping of reconstructed lineage-specific rediploidization histories contrasting Ssa06 from Atlantic salmon with its orthologous region in brown trout (Str01) demonstrating the persistence of intermixing of phylogenetic signals (i.e. different rediploidization histories) in both genome assemblies. All other details are as described in the Supplementary Fig. 2 legend.

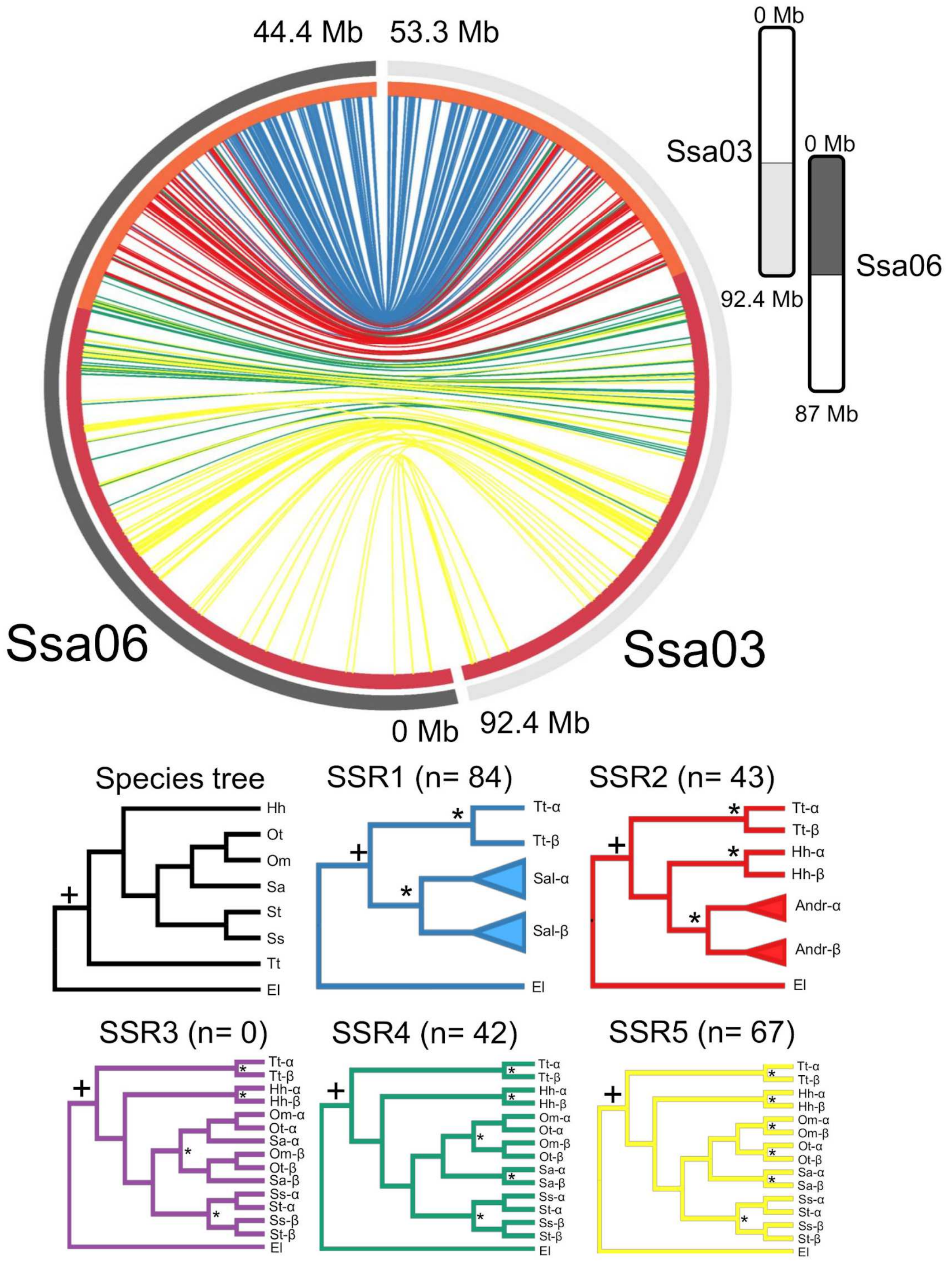


**Supplementary Fig. 13.** Mapping of reconstructed lineage-specific rediploidization histories for ohnologue regions Ssa03 and Ssa06 in the Atlantic salmon genome, using protein-coding gene trees as opposed to our genome-wide alignments.

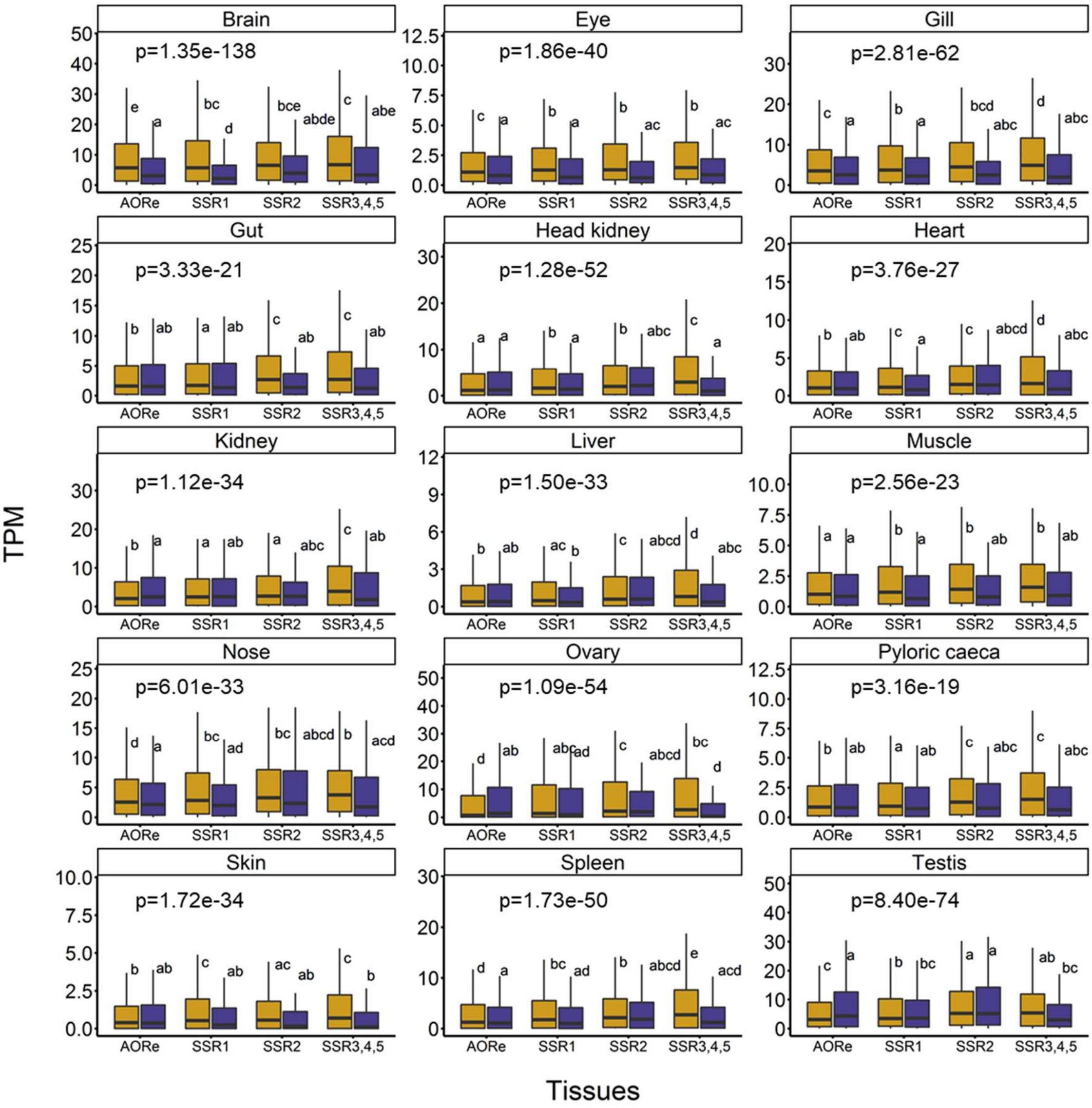


**Supplementary Fig. 14.** Gene expression of ohnologue (yellow shading) and singleton gene (blue shading) sets residing in genomic regions with distinct rediploidization ages. RNA-Seq transcripts per million (TPM) data for Atlantic salmon tissues from Lien et al. 2016. Box and whisker plot definition: each box spans the interquartile range, with the median (Q2) as a central bar, and upper and lower bounds representing the respective minimum and maximum values within the 25th percentile (Q1) and 75th percentile (Q3). The upper and lower whisker bounds represent the largest and smallest values lying within 1.5 times above Q3 and below Q1, respectively. One-way ANOVA was performed to test for differences in TPM across the shown gene sets (overall *P* value given in graphs). Tukey's test was performed to identify category specific TPM differences (differences among categories at 95% confidence level shown by different letters).

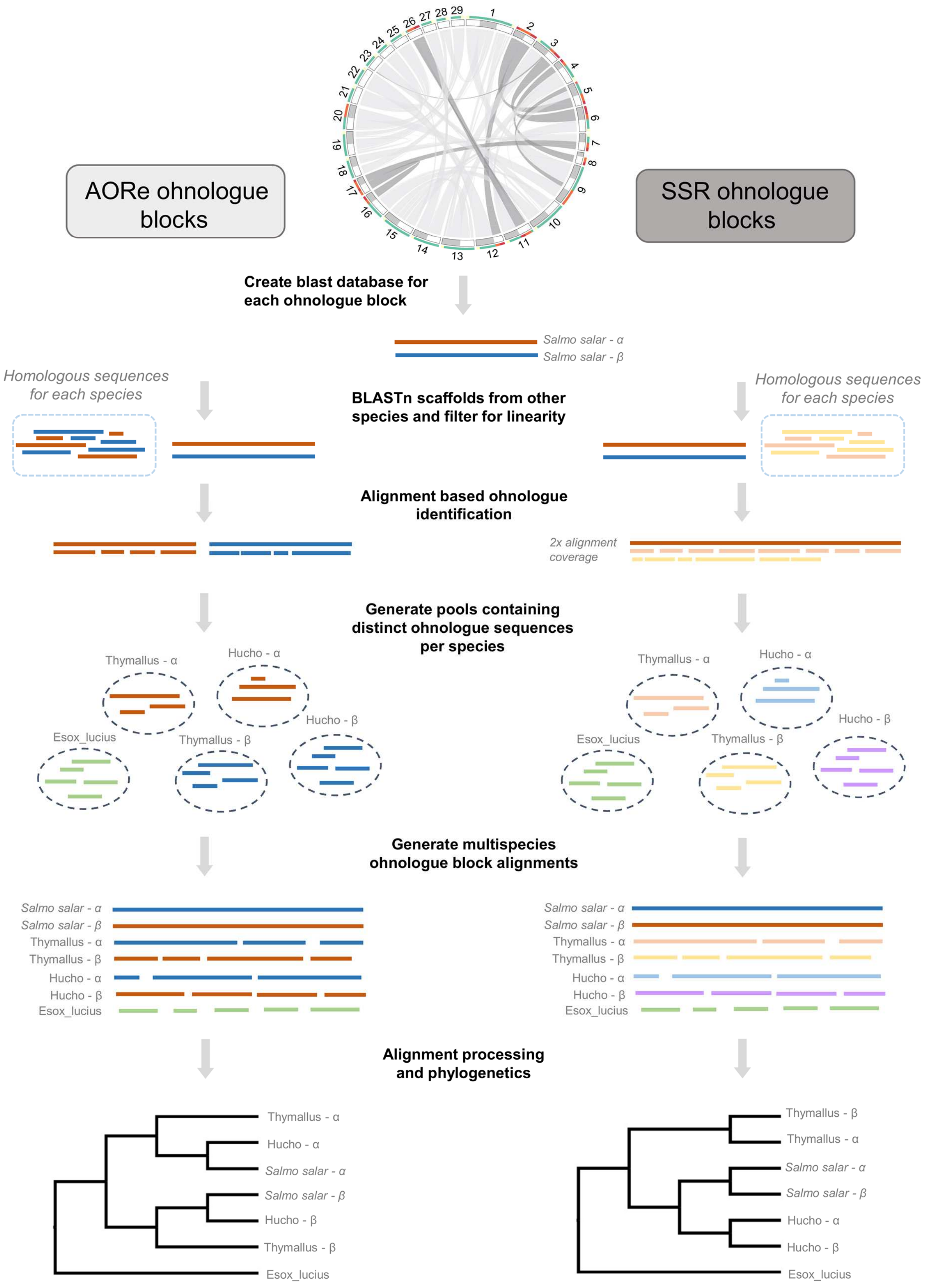


**Supplementary Fig. 15.** Summary of steps used to generate multispecies ohnologue alignments in advance of phylogenomic analyses.

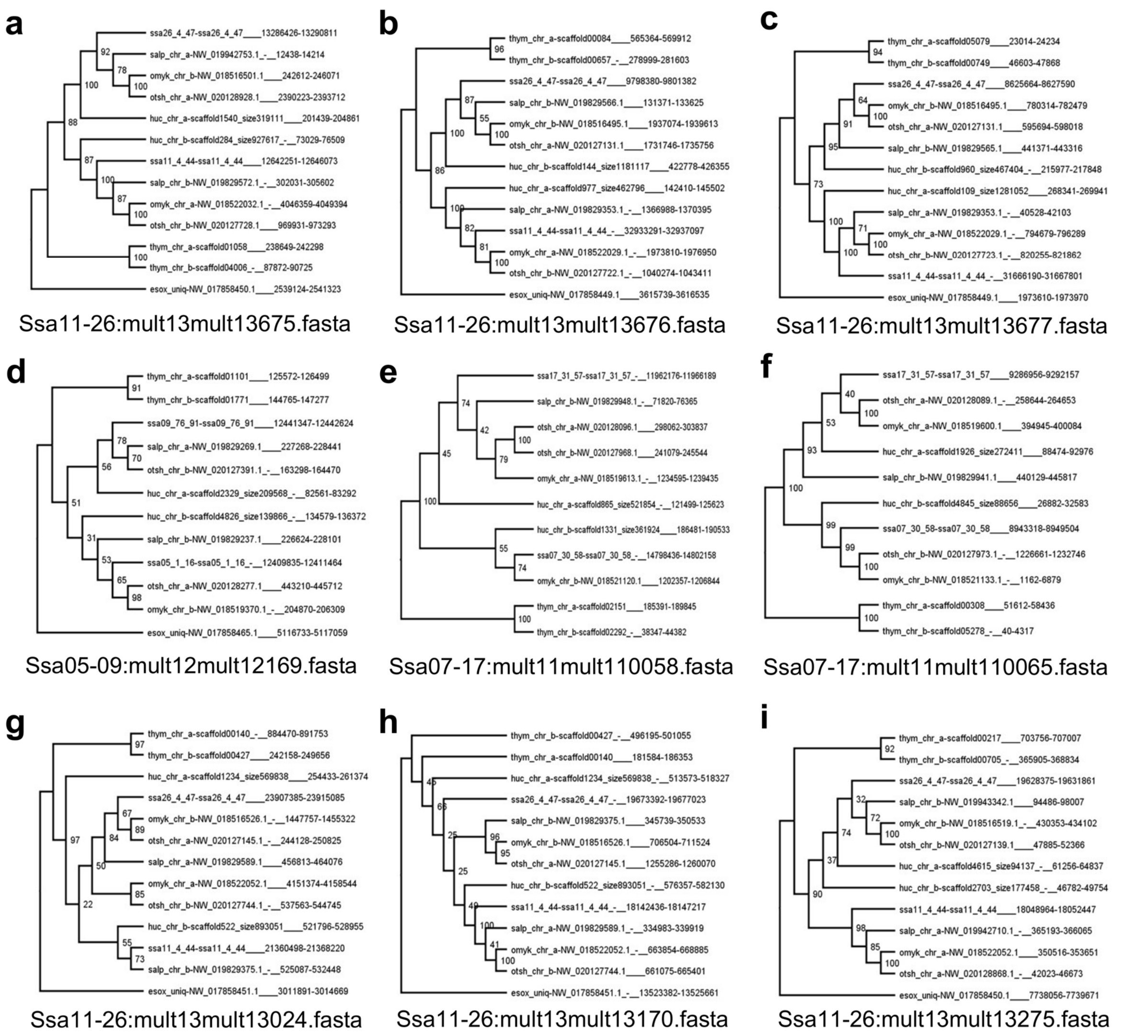


**Supplementary Fig. 16.** Example trees for SSR1 topology. **a-c**. Example trees matching exactly to the predicted topology with no ambiguity, **d-f**. Example trees with minor ambiguity in branching but overwhelmingly supporting the expected topology, **g-i**. Example trees rejected due to ambiguous topology or low support. The numbers at the nodes of each tree indicate bootstrap support values Species abbreviations: thym = European grayling; ssa = Atlantic salmon; salp = Arctic charr; omyk = rainbow trout; otsh = chinook salmon; huc = huchen; esox = northern pike. All SSR trees are provided in Supplementary Data 7 and their genomic coordinates in Atlantic salmon are given in Supplementary Data 8.

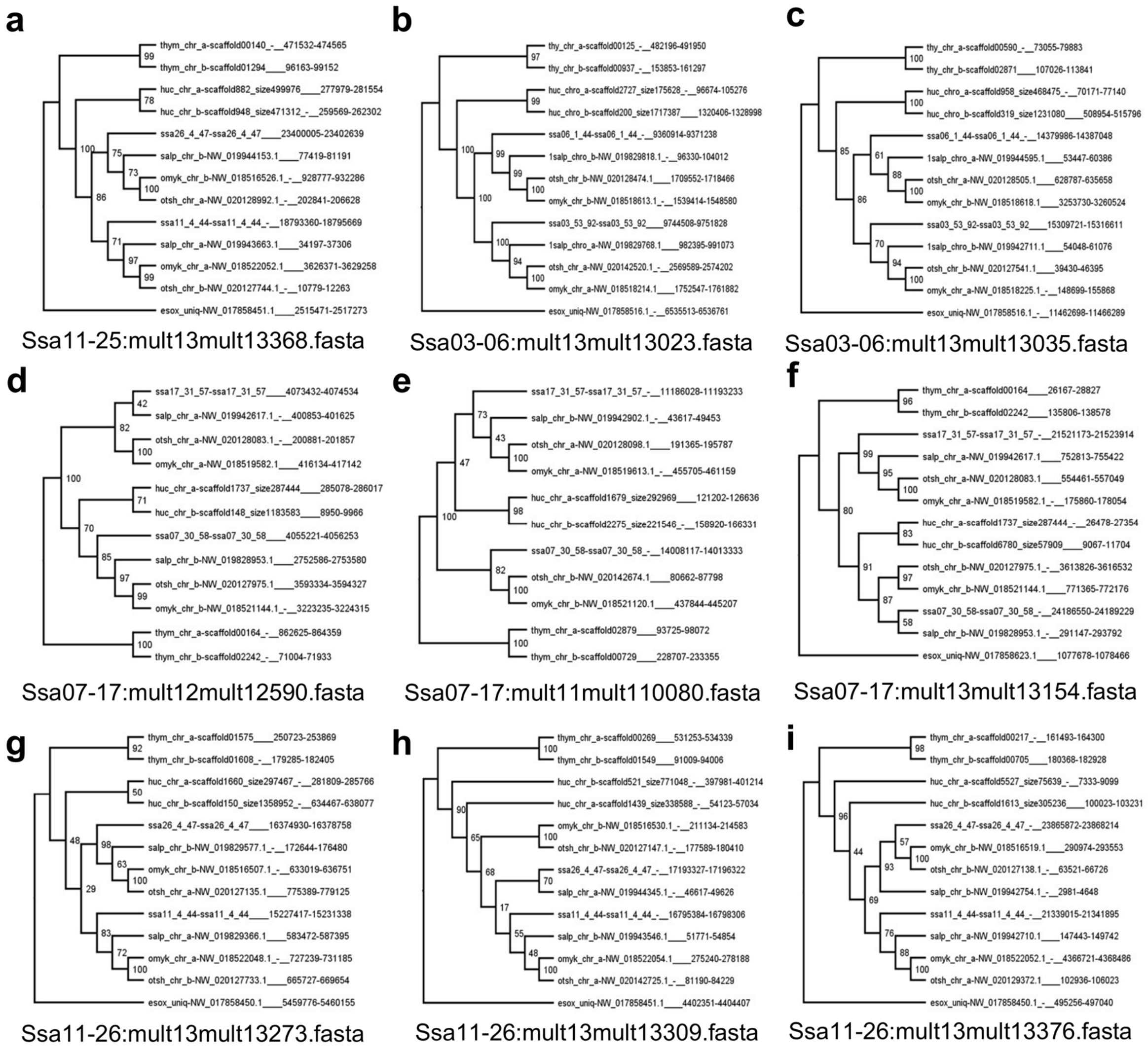


**Supplementary Fig. 17.** Example trees for SSR2 topology. Other details as in the Supplementary Fig. 16 legend.

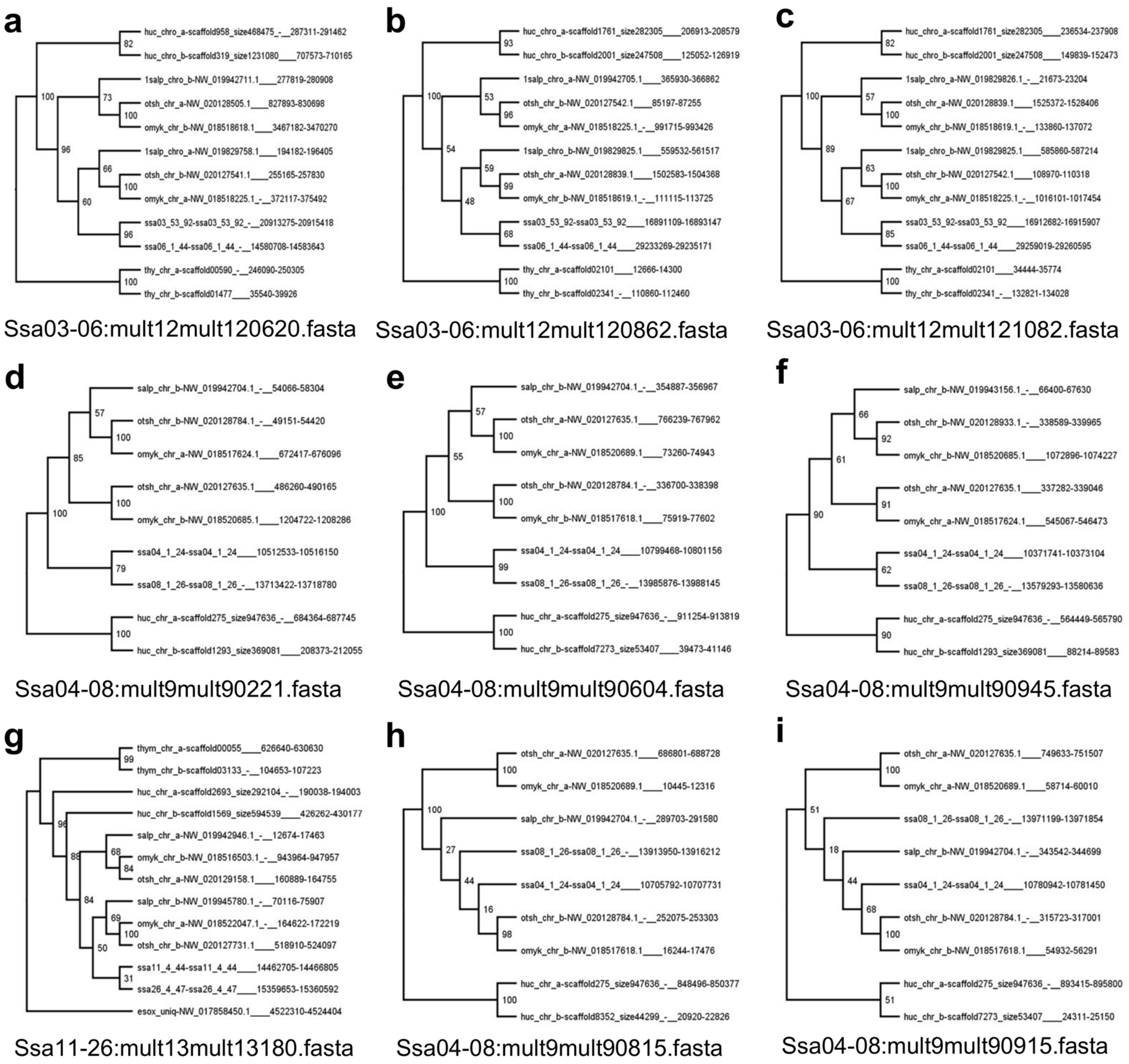


**Supplementary Fig. 18.** Example trees for SSR3 topology. Other details as in the Supplementary Fig. 16 legend.

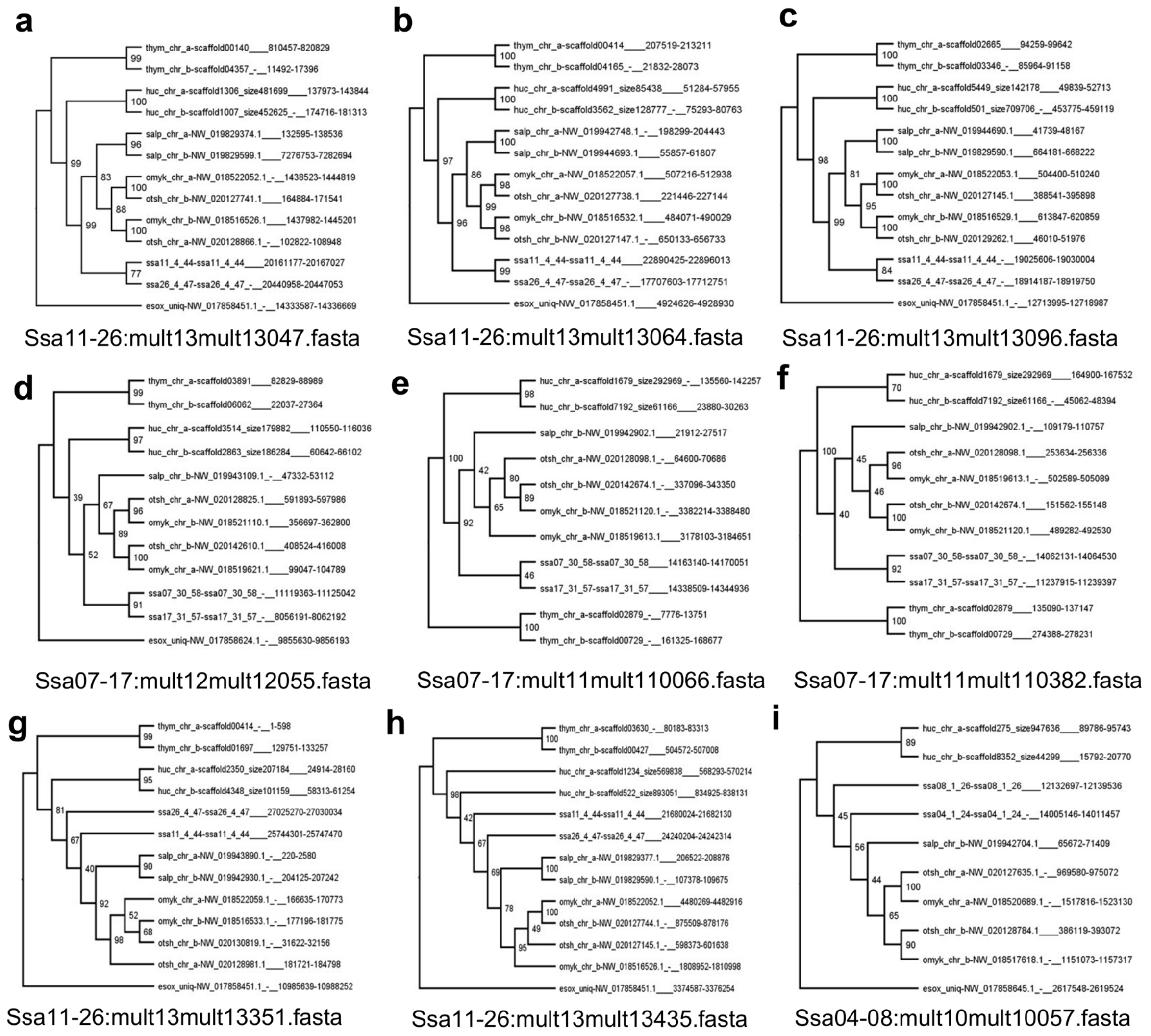


**Supplementary Fig. 19.** Example trees for SSR4 topology. Other details as in the Supplementary Fig. 16 legend.

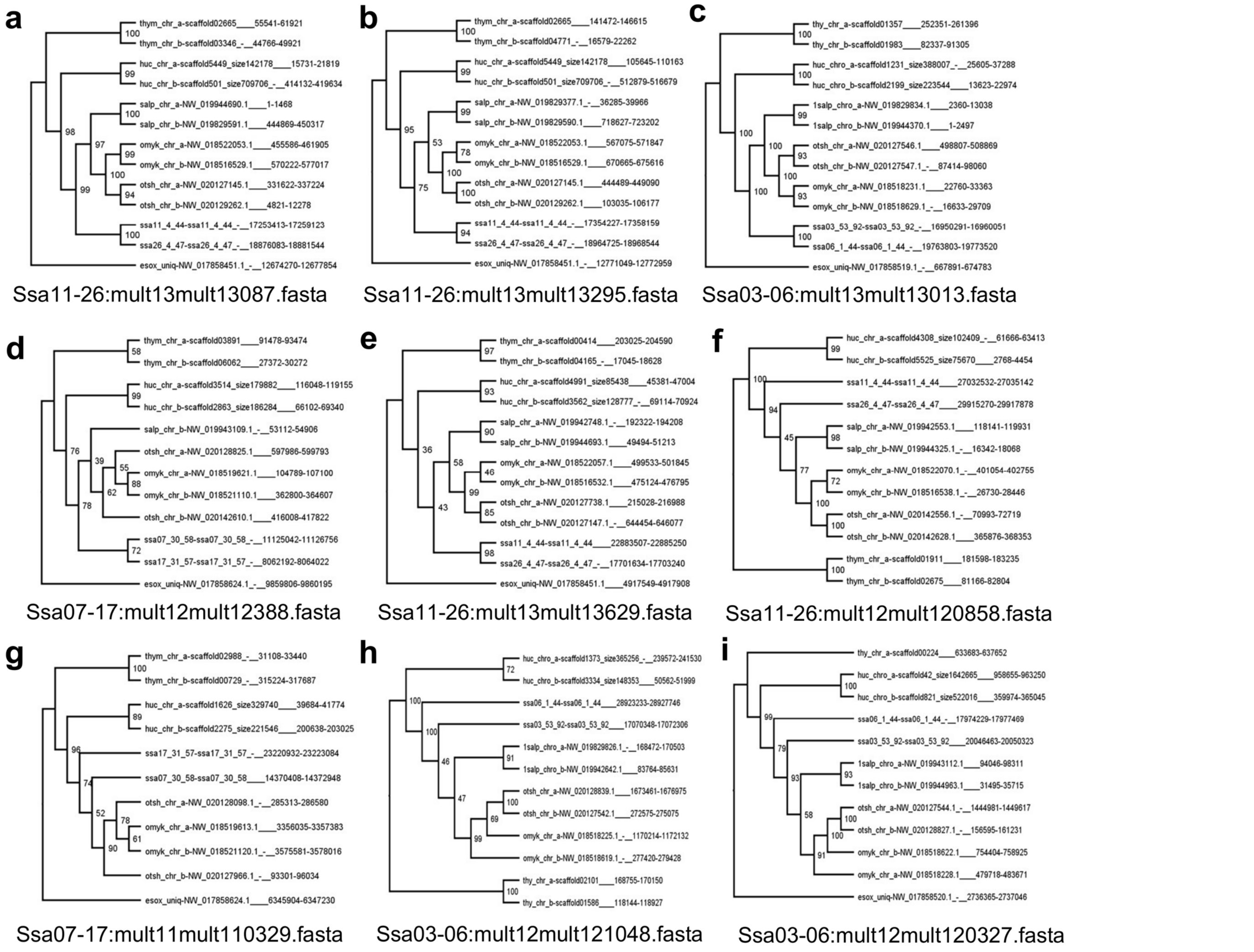


**Supplementary Fig. 20.** Example trees for SSR5 topology. Other details as in the Supplementary Fig. 16 legend.

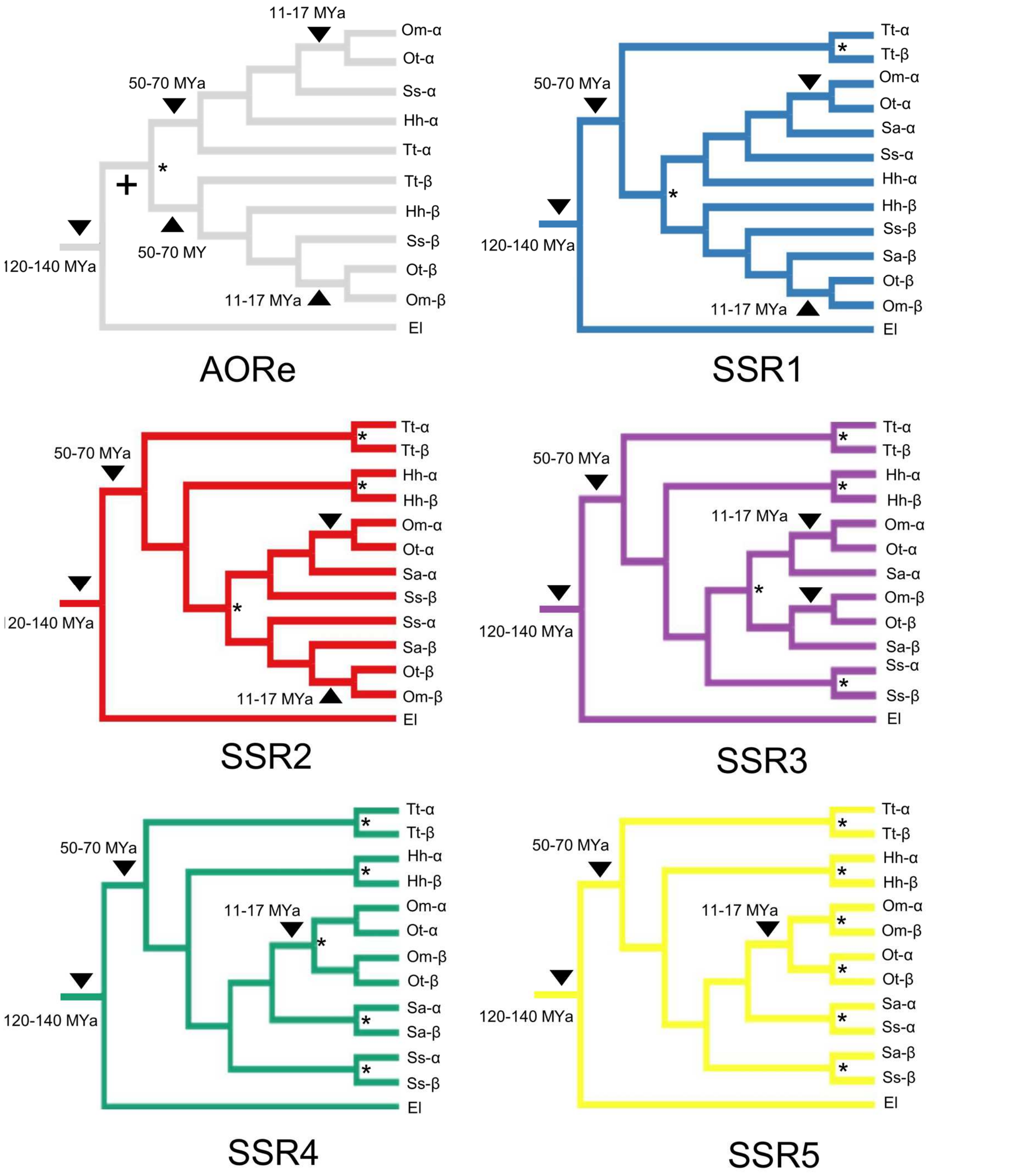


**Supplementary Fig. 21.** Temporal constraints used in Bayesian rediploidization age analysis for genomic regions with distinct rediploidization histories. The shown topologies were set in the MCMCtree analyses, with black triangles indicating the position of temporal calibrations, which were set as uniform distribution priors.

#### Supplementary Tables

**Supplementary Table 1.** Statistics for huchen genome sequencing data

| Data type | Library | Read pairs | Total data (Gb) | [illegible]overage | Mean read length (bp) |
|---|---|---|---|---|---|
| Raw sequencing data | *Paired-end* | 397,529,369 | 198.76 | 90.34 | 250 |
| | *6Kb mate-pair* | 78,491,938 | 39.24 | 17.83 | 250 |
| | 12Kb mate-pair | 213,236,078 | 106.61 | 48.46 | 250 |
| | Total: | 689,257,385 | 344.61 | [illegible]56.63 | 250 |
| Post quality control | Paired-end | 370,394,584 | 139.26 | 63.3 | 188 |
| | 6Kb mate-pair | 44,935,049 | 11.68 | 5.31 | 131 |
| | 12Kb mate-pair | 124,687,467 | 32.29 | 14.70 | 129 |
| | Total: | 520,017,100 | 178.17 | 83.31 | N/A |

**Supplementary Table 2**. Statistics for the final huchen genome assembly

| Statistic | All sequences |
|---|---|
| Assembly Size | 2,487,549,814 |
| Ungapped length | 1,917,049,985 |
| Scaffold N50 | 287,338 Kb |
| Contig N50 | 37,369 |
| Longest Scaffold | 4.603 Mb |
| No scaffolds | 71,639 |
| % GC content | 42.61 % |
| Complete BUSCOs | 90.2% |
| Complete – single copy | 48.6 % |
| Complete - duplicated | 41.6 % |
| Fragmented | 3.2 % |
| Missing | 6.6% |
| CEGMA-Complete | 236 |
| CEGMA-Fragmented | 4 |
| CEGMA-Missing | 8 |

**Supplementary Table 3**. Repeat prediction within the huchen genome

| Element | | Number of elements | length | Percentage |
|---|---|---|---|---|
| SINEs | | 20,347 | 4,143,480 | 0.17 |
| | AIUs | 1 | 64 | 0 |
| | MIRs | 0 | 0 | 0 |
| LINEs | | 202,026 | 100,766,745 | 4.05 |
| | LINE1 | 3,809 | 1,831,205 | 0.07 |
| | LINE2 | 71,383 | 410,39,218 | 1.65 |
| | L3/CR1 | 2,012 | 877,215 | 0.04 |
| LTR elements | | 58,047 | 36,317,611 | 1.46 |
| | ERVL | 3 | 321 | 0 |
| | ERVL-MaLRs | 0 | 0 | 0 |
| | ERV_classI | 7,818 | 5,592,413 | 0.22 |
| | ERV_classII | 162 | 138,073 | 0.01 |
| DNA elements | | 904,622 | 308,552,003 | 12.40 |
| | hAT-Charlie | 7,988 | 2,777,233 | 0.11 |
| | TcMar-Tigger | 70 | 31,297 | 0 |
| Unclassified | | 708,405 | 428,579,333 | 17.23 |
| Total interspersed repeats | | | 878,359,172 | 35.31 |
| Small RNA | | 75 | 9,505 | 0 |
| Satellites | | 107,39 | 4,887,842 | 0.2 |
| Simple repeats | | 661,233 | 44,896,670 | 1.8 |
| Low complexity | | 94,780 | 7,661,339 | 0.31 |
| TOTAL | | | 932,098,879 | 37.47 |
